# CCR5 deficiency impairs CD4^+^ T cell memory responses and antigenic sensitivity through increased ceramide synthesis

**DOI:** 10.1101/2020.02.14.948893

**Authors:** Ana Martín-Leal, Raquel Blanco, Josefina Casas, María E. Sáez, Elena Rodríguez-Bovolenta, Itziar de Rojas, Carina Drechsler, Luis Miguel Real, Gemma Fabrias, Agustín Ruíz, Mario Castro, Wolfgang W.A. Schamel, Balbino Alarcón, Hisse M. van Santen, Santos Mañes

**Author notes:** Correspondence to: Santos Mañes, Centro Nacional de Biotecnología/CSIC, Department of Immunology and Oncology, Darwin 3, 28049 Madrid, Spain. These authors contributed equally to this work (Ana Martín-Leal, Raquel Blanco).

## Abstract

In CD4^+^ T cells, CCR5 is not only a coreceptor for HIV-1 infection, but also contributes to their functional fitness. Here we show that by limiting GATA-1-induced transcription of specific ceramide synthases, CCR5 signaling reduces ceramide levels and thereby increases T cell antigen receptor (TCR) nanoclustering in antigen-experienced mouse and human CD4^+^ T cells. This activity is CCR5-specific and independent of CCR5 costimulatory activity. CCR5-deficient mice showed reduced production of high affinity class-switched antibodies, but only after antigen rechallenge, which implies an impaired memory CD4^+^ T cell response. This study identifies a CCR5 function in the generation of CD4^+^ T cell memory responses, and establishes an antigen-independent mechanism that regulates TCR nanoclustering by altering specific lipid species.

## INTRODUCTION

The C-C motif chemokine receptor 5 (CCR5) is a seven-transmembrane G protein-coupled receptor (GPCR) expressed on the surface of several innate and adaptive immune cell subtypes, including effector and memory CD4^+^ T lymphocytes (Gonzalez-Martin et al., 2012). CCR5 acts also a necessary coreceptor for infection by HIV-1. An HIV-resistant population served to identify a 32-bp deletion within the CCR5 coding region (*ccr5*Δ32), which yields a non-functional receptor (Blanpain et al., 2002). Since *ccr5*Δ32 homozygous individuals are seemingly healthy, a radical body of thought considers that CCR5 is dispensable for immune cell function.

Experimental and epidemiological evidence nonetheless indicates that CCR5 has an important role in innate and acquired immune responses. CCR5 and its ligands C-C motif ligand 3 [CCL3; also termed macrophage inflammatory protein (MIP)-1α], CCL4 (MIP-1β), CCL5 [regulated upon activation, normal T cell expressed and secreted (RANTES)] and CCL3L1 have been associated with exacerbation of chronic inflammatory and autoimmune diseases. Despite varying information due probably to ethnicity effects (Lee et al., 2013, Schauren et al., 2013), further complicated in admixed populations (Toson et al., 2017), epidemiological studies support the *ccr5*Δ32 allele as a marker for good prognosis for these overreactive immune diseases (Vangelista & Vento, 2017). In contrast, *ccr5*Δ32 homozygotes are prone to fatal infections by several pathogens such as influenza, West Nile, and tick-borne encephalitis viruses (Ellwanger & Chies, 2019, Falcon et al., 2015, Lim & Murphy, 2011). The mechanisms by which the *ccr5*Δ32 polymorphism affects all these pathologies have usually been linked to the capacity of CCR5 to regulate leukocyte trafficking. For example, CCR5 deficiency reduces recruitment of influenza-specific memory CD8^+^ T cells and accelerates macrophage accumulation in lung airways during virus rechallenge (Dawson et al., 2000, Kohlmeier et al., 2008); this could lead to acute severe pneumonitis, a fatal flu complication. CCR5 nonetheless has migration-independent functions that maximize T cell activation by affecting immunological synapse (IS) formation (Floto et al., 2006, Franciszkiewicz et al., 2009, Molon et al., 2005) as well as T cell transcription programs associated with cytokine production (Camargo et al., 2009, Lillard et al., 2001). CCR5 and its ligands are also critical for cell-mediated immunity to tumors and pathogens, including HIV-1 (Bedognetti et al., 2013, Dolan et al., 2007, González-Martín et al., 2011, Ugurel et al., 2008).

Whereas the role of CCR5 in T cell priming is well-established, its involvement in memory responses has not been addressed in depth. Only a single report suggested CCR5 involvement in CD4^+^ T cell promotion of memory CD8^+^ T cell generation through a migration-dependent process (Castellino et al., 2006). It remains unknown whether CCR5 endows memory T cells with additional properties. One such property is the elevated sensitivity of effector and memory (“antigen-experienced”) CD4^+^ and CD8^+^ T cells to their cognate antigen compared to naïve cells (Huang et al., 2013, Kersh et al., 2003, Kimachi et al., 1997). This sensitivity gradient (memory>>effector>naïve) in CD8^+^ T cells is linked to increased valency of preformed TCR oligomers at the cell surface, termed TCR nanoclusters (Kumar et al., 2011). This antigen-independent TCR nanoclustering (Lillemeier et al., 2010, Schamel & Alarcon, 2013, Schamel et al., 2005, Schamel et al., 2006, Sherman et al., 2011) enhances antigenic sensitivity by increasing avidity to multimeric peptide-major histocompatibility complexes (Kumar et al., 2011, Molnar et al., 2012) and by allowing cooperativity between TCR molecules (Martín-Blanco et al., 2018, Martínez-Martín et al., 2009). TCRβ subunit interaction with cholesterol (Chol) and the presence of sphingomyelins (SM) are both essential for TCR nanoclustering (Beck-Garcia et al., 2015, Molnar et al., 2012). Replacement of Chol by Chol sulfate impedes TCR nanocluster formation and reduces CD4^+^CD8^+^ thymocyte sensitivity to weak antigenic peptides (Wang et al., 2016). Whether antigen-experienced CD4^+^ T cell sensitivity is linked to TCR nanoscopic organization and the homeostatic factors that regulate TCR nanoclustering remains unexplored.

Given its costimulatory role in CD4^+^ T cells, we speculated that CCR5 signals would affect the antigenic sensitivity of CD4^+^ memory T cells. To test this hypothesis, we analyzed the function of *in vivo*-generated memory CD4^+^ T cells in wild-type (WT) and CCR5^−/−^ mice, and the effect of CCR5 deficiency on CD4 T cell help in the T-dependent humoral response. We found that CCR5 is necessary for the establishment of a functional CD4 memory response through a mechanism independent of its co-stimulatory role for the TCR signal. We show that CCR5 deficiency does not affect memory CD4 T cell generation, but reduces their sensitivity to antigen. Our data demonstrate an unreported CCR5 regulatory role in memory CD4^+^ T cell function by inhibiting the synthesis of ceramides, which are identified here as negative membrane regulators of TCR nanoscopic organization.

## RESULTS

### CCR5 deficiency impairs the CD4^+^ T cell memory response

To determine the role of CCR5 in CD4^+^ memory T cell generation and/or function, we adoptively transferred congenic CD45.1 mice with lymph node/spleen cell suspensions from OT-II WT or CCR5^−/−^ mice (CD45.2), and subsequently infected them with OVA-encoding vaccinia virus; five weeks post-immunization, we analyzed spleen CD45.2^+^ donor cells from OT-II mice. CCR5 expression on OT-II cells affected neither the total number of memory CD4^+^ T cells (Fig.1A, B) nor the percentage of CD4^+^ T_EM_ (CD44^hi^; CD62L^−^; Fig.1C) or T_CM_ (CD44^hi^; CD62L^+^; Fig. 1D) cells generated. OT-II WT cells nonetheless had stronger responses to antigenic restimulation than OT-II CCR5^−/−^ memory T cells, as determined by the percentage of interferon (IFN)γ-producing cells after *ex vivo* stimulation with OVA_323-339_ (Fig.1E).

**Figure 1.**
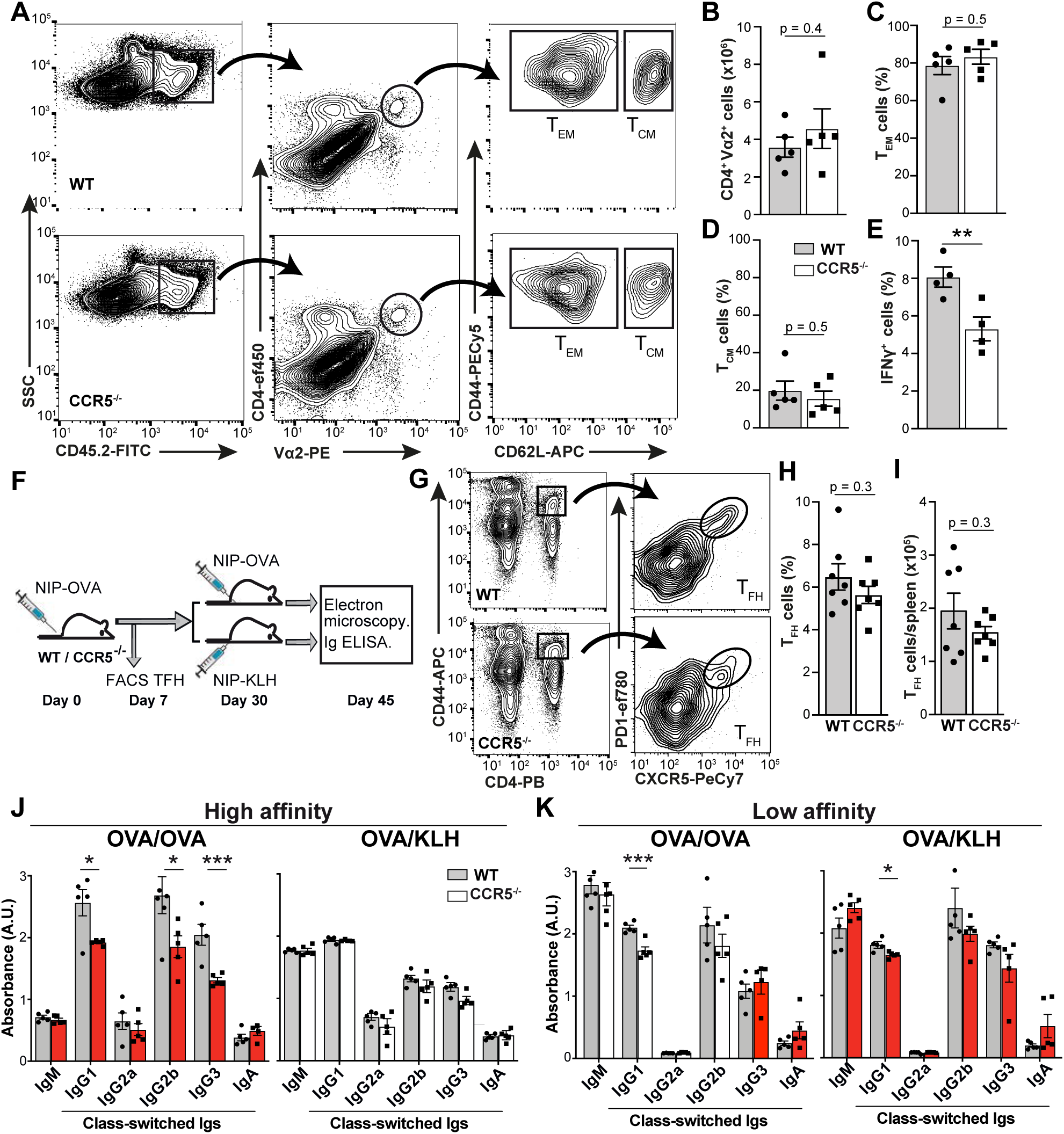
CCR5 deficiency impairs CD4^+^ T cell memory responses. **A.** Representative plots of splenocytes from CD45.1 mice adoptively transferred with CD45.2 OT-II WT or CCR5^−/−^ lymph node cell suspensions, five weeks after infection with rVACV-OVA virus. The gating strategy used to identify the memory CD4^+^ T cell subtypes is shown (*n* = 5). **B.** Absolute number of OT-II cells recovered in spleens of mice as in *a*. **C, D.** Percentage of CD4^+^ T_EM_ (*C*) and T_CM_ (*D*) in the OT-II WT and CCR5^−/−^ populations. **E.** IFNγ-producing OT-II WT and CCR5^−/−^ memory cells isolated from mice as in *a* and restimulated *ex vivo* with OVA_323-339_ (1 μM). **F.** Immunization scheme for NIP-OVA and NIP-KLH in WT and CCR5^−/−^ mice. **G-I.** Representative plots (*G*) and quantification of the frequency (*H*) and absolute number (*I*) of T_fh_ cells (CD4^+^CD44^+^PD-1^+^CXCR5^+^) in the spleen after primary immunization (day 7) with NIP-OVA. **J, K.** ELISA analysis of high- (*J*) and low-affinity (*K*) isotype-specific anti-NIP antibodies in sera from OVA/OVA- and OVA/KLH-immunized mice (day 15 post-challenge; *n* = 5 mice/group). Data representative of one experiment of two. *B-E, H-K*, data are mean ± SEM. * p <0.05, ** p <001, *** p <0.001, two-tailed unpaired Student’s *t*-test.

We also studied T cell-dependent B cell responses in WT and CCR5^−/−^ mice after immunization with the hapten 4-hydroxy-3-iodo-5-nitrophenylacetyl coupled to ovalbumin (NIP-OVA; Fig.1F). We detected no difference in the percentage or absolute number of T follicular helper (T_fh_) cells (CD4^+^, CD44^hi^, CXCR5^+^, PD1^+^) between WT and CCR5^−/−^ mice at seven days post-immunization (Fig. 1G-I). At day 30, half of the mice were boosted with the same NIP-OVA immunogen (OVA/OVA) and the other half received NIP conjugated with another carrier protein (OVA/KLH); levels of NIP-specific high and low affinity immunoglobulins (Ig) were analyzed 15 days later. Comparison of the humoral responses between OVA/OVA- and OVA/KLH-immunized mice would assess the effect of memory CD4^+^ T cells specific for the first carrier protein on the humoral response to NIP. There were no differences in high/low affinity NIP-specific IgM production between WT and CCR5^−/−^ mice with either immunization strategy (Fig. 1J, K). CCR5 deficiency markedly impaired the generation of high-affinity class-switched anti-NIP antibodies specifically in OVA/OVA-immunized mice (Fig. 1J, K). Since class switching was similar in WT and CCR5^−/−^ OVA/KLH-immunized mice, our results suggest that CCR5 deficiency reduces the generation of high-affinity class-switched immunoglobulins due to deficient memory CD4^+^ T cell function.

### The CCR5 effect on antigen-experienced CD4^+^ T cells is cell-autonomous

To test whether the *in vivo* memory defect associated to CCR5 deficiency was intrinsic to CD4^+^ T cells, we activated OT-II WT and CCR5^−/−^ spleen T cells with OVA_323-339_ antigen for three days; after antigen removal, we cultured cells for four days with IL-2 or IL-15. OT-II cells that differentiated in exogenous IL-2 expressed CCL3, CCL4, CCL5, and a functional CCR5 receptor, as determined by their ability to migrate in a CCL4 gradient (Fig. S1A-C).

Like CD8^+^ T cells (Richer et al., 2015), OT-II cells cultured with IL-15 showed a memory-like phenotype; they were smaller than IL-2-cultured cells and retained CD62L with reduced activation marker expression (CD25, CD69, CD44) compared to IL-2-cultured T cells (Fig. 2A). Findings were similar in OT-II WT and CCR5^−/−^ cells (Fig. 2B), which reinforced the idea that CCR5 is not involved in CD4^+^ T memory cell differentiation. Restimulation of IL-2- or IL-15-expanded OT-II lymphoblasts with the OVA_323-339_ peptide nonetheless indicated that CCR5-expressing cells showed strong proliferation and higher IL-2 production at low antigen concentrations than CCR5-deficient cells (Fig. 2C-F), indicative of an increased number of cells responding to antigenic stimulation. CCR5 might thus increase the antigenic sensitivity of antigen-experienced CD4^+^ T cells in a cell-autonomous manner.

**Figure 2.**
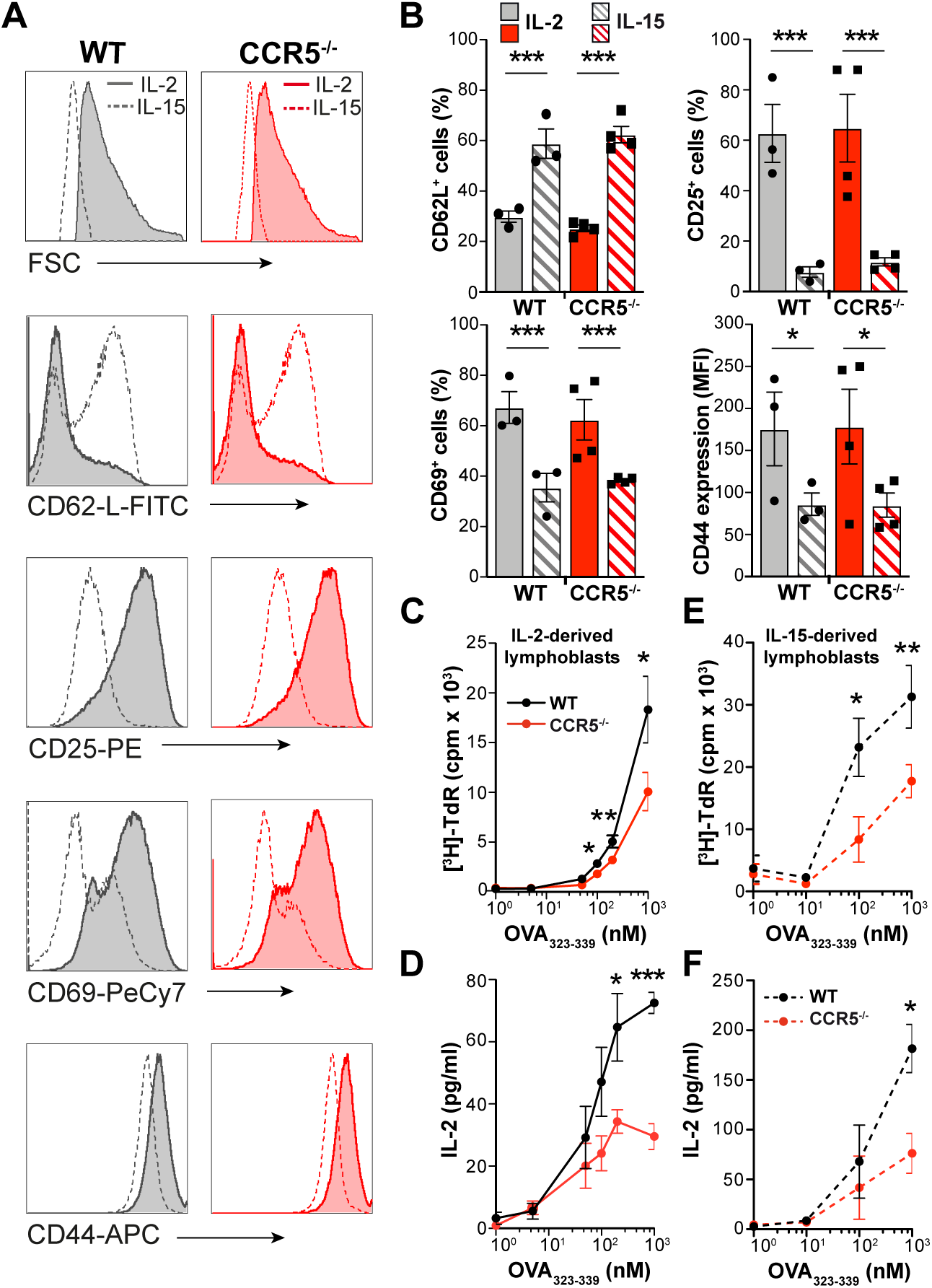
CCR5 increases the sensitivity of antigen-experienced CD4^+^ T cells. **A, B.** Representative histograms (*A*) and quantification of mean fluorescence intensity (MFI) or the percentage of cells positive for the indicated memory markers (*B)* in OT-II WT and CCR5^−/−^ lymphoblasts expanded in IL-2 or IL-15, as specified. Data shown as mean ± SEM (*n* ≥3). **C-F.** IL-2- (*C*, *D*) and IL-15-expanded lymphoblasts (*E*, *F*) were restimulated with indicated concentrations of OVA_323-339_; cell proliferation (thymidine incorporation into DNA; *C*, *E*) and IL-2 production (by ELISA; *D*, *F*) were measured after 72 h. Data shown as mean ± SEM (*n* = 5). * p <0.05, ** p <001, *** p <0.001, two-way ANOVA (*B*) or two-tailed unpaired Student’s *t*-test (*C-F*).

### CCR5 modulates TCR nanoclustering in antigen-experienced CD4^+^ T cells

The high antigenic sensitivity of antigen-experienced CD8^+^ T cells was partially attributed to increased TCR nanoclustering (Kumar et al., 2011). To determine whether CCR5 deficiency influences TCR organization, we used electron microscopy (EM) to analyze surface replicas of OT-II WT and CCR5^−/−^ naïve cells and lymphoblasts after labeling with anti-CD3ε antibody and 10 nm gold-conjugated protein A. We found no differences in TCR nanoclusters between OT-II WT or CCR5^−/−^ naïve cells, which had a small percentage of TCR nanoclusters larger than 4 TCR in both genotypes (Fig. 3A). In contrast, there was a significant increase in TCR nanocluster number and size in WT compared to CCR5^−/−^ lymphoblasts (Fig. 3B, C). As predicted, there was a gradient in TCR nanoclustering of naïve << IL-2-< IL-15-differentiated OT-II WT cells (Fig. S1D), which coincided with increased antigenic sensitivity of the IL-15-expanded cells (Fig. S1E). These findings thus reinforce the IL-15-induced memory-like phenotype versus the IL-2-induced effector-like phenotype, and link TCR nanoclustering with increased sensitivity in antigen-experienced CD4^+^ T cells. The difference in TCR nanoclustering between WT and CCR5^−/−^ cells was nevertheless similar in IL-2- and IL-15-expanded lymphoblasts, which indicates that CCR5 affects TCR nanoclustering in lymphoblasts independently of the cytokine milieu.

**Figure 3.**
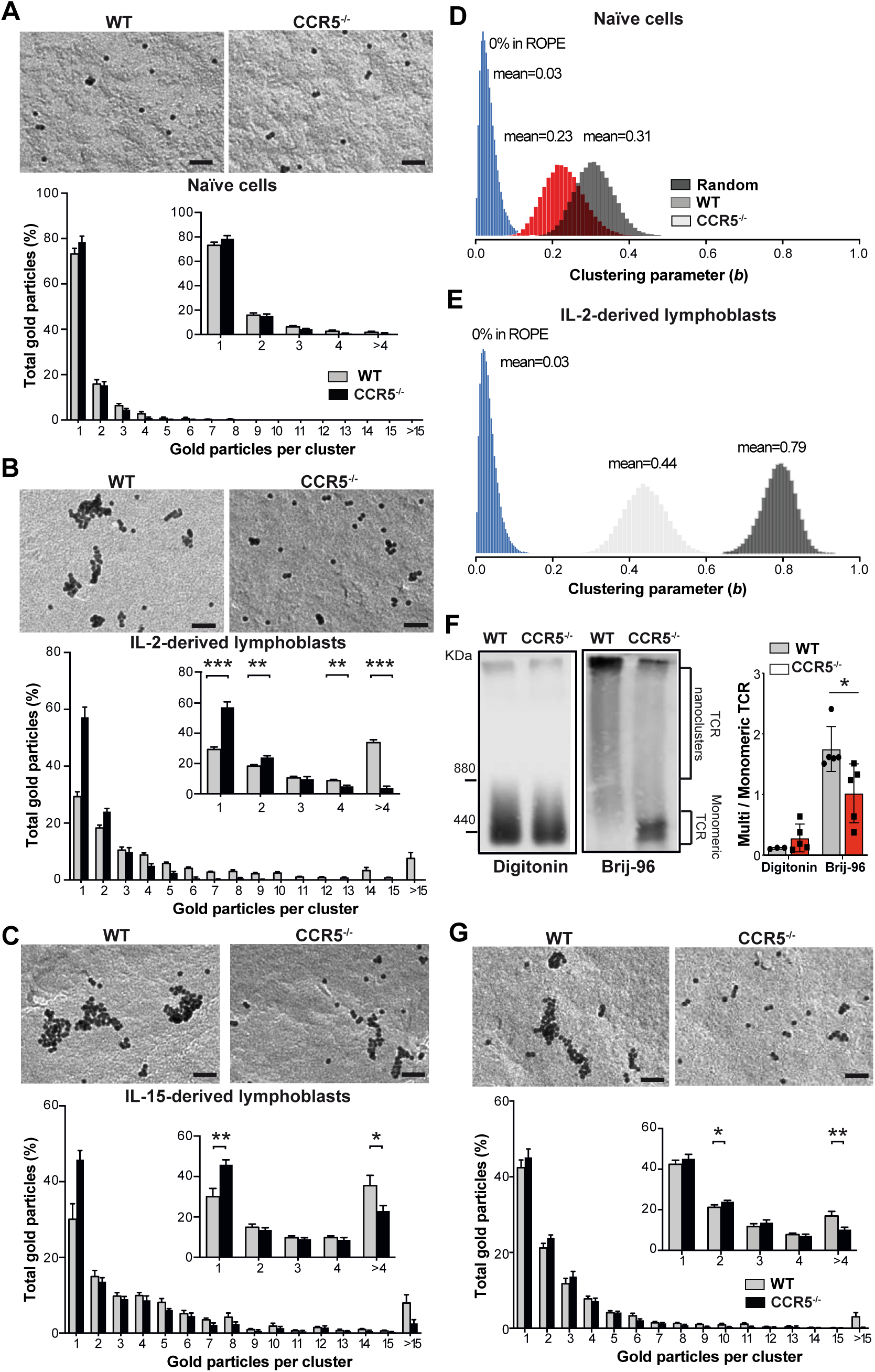
CCR5 increases TCR nanoclustering in antigen-experienced CD4^+^ T cells. **A-C.** Analysis of TCR nanoclustering in OT-II WT and CCR5^−/−^ naïve cells (*A*; *n* = 6 cells/genotype; WT: 3427, CCR5^−/−^: 3528 particles), and IL-2- (*B*; WT, *n* = 8 cells, 15419 particles; CCR5^−/−^, *n* = 6 cells, 5410 particles) or IL-15-expanded lymphoblasts (*C*; WT, *n* = 8 cells, 27518 particles; CCR5^−/−^, *n* = 7 cells, 22696 particles). A representative small field image at the top of each panel shows gold particle distribution in the cell surface replicas of anti-CD3ε-labeled cells (bar, 50 nm); at bottom, quantification (mean ± SEM) of gold particles in clusters of indicated size in WT (gray bars) and CCR5^−/−^ cells (black). Insets show the distribution between clusters of one, two, three, four, or more than four particles, and statistical analysis (* p <0.05, ** p <001, *** p <0.001, one-tailed unpaired Student’s *t*-test). **D, E.** Posterior distribution in naïve (*D*) and IL-2-expanded lymphoblasts (*E*) of the clustering parameter *b* for WT (gray) and CCR5^−/−^ cells (red); randomly generated distributions of receptors are shown in blue. The mean value of the *b* parameter is indicated for each condition. The probability of a chance distribution similar to that determined in cells is nearly 0% by the ROPE. **F.** Comparison of TCR oligomer size using BN-PAGE and anti-CD3ζ immunoblotting in day 10, IL-2-expanded WT and CCR5^−/−^ OT-II lymphoblasts lysed in buffer containing digitonin or Brij-96. The marker protein is ferritin (f1, 440 and f2, 880 kDa forms). The ratio of TCR nanoclusters to monomeric TCR in each lysis condition was quantified by densitometry (bottom). Data shown as mean ± SEM (*n* = 5). **G.** Top, representative small field images showing gold particle distribution in the cell surface replicas of CD4^+^ T cells isolated from OVA/OVA-immunized WT and CCR5^−/−^ mice. Bottom, quantification (mean ± SEM) of gold particles in clusters of the indicated size (WT, gray bars; *n* = 5 cells, 14680 particles; CCR5^−/−^, black; *n* = 7 cells, 15374 particles). Insets show the distribution between clusters of one, two, three, four or more than four particles, and statistical analysis. * p <0.05, ** p <001, *** p <0.001, one-tailed unpaired Student’s *t*-test. Bar, 50 nm.

Using a Monte-Carlo simulation, we applied data from surface replicas of naïve and IL-2-expanded OT-II lymphoblasts to determine whether the experimental frequency of cluster size was due to random distribution of gold particles. In all cases, the cluster distributions observed experimentally differed significantly from pure random proximity between clusters (Fig. S2). To define the differences between OT-II WT and CCR5^−/−^ cells, we used a model that accounts for receptor clustering dynamics (Castro et al., 2014), a Bayesian inference method that estimates the so-called clustering parameter, *b*. Based on this model, we concluded that the probability of a chance nanocluster distribution similar to that observed for naïve and activated OT-II WT and CCR5^−/−^ cells approaches 0% (Fig. 3D, E). Posterior distribution analysis also showed that whereas the clustering parameter was very similar between naïve OT-II WT and CCR5^−/−^ cells (Fig. 3D), there was clear separation in lymphoblasts (Fig. 3E). These analyses provide a mathematical framework that validates the TCR nanoclustering differences between WT and CCR5^−/−^ cells, as determined by EM.

The differences in TCR oligomerization between OTII WT and CCR5^−/−^ lymphoblasts were also studied using blue-native gel electrophoresis (BN-PAGE) (Schamel et al., 2005, Swamy & Schamel, 2009). Cell lysis with digitonin, a detergent that disrupts TCR nanoclusters into their monomeric components, showed that WT and CCR5^−/−^ lymphoblasts expressed comparable TCR levels, as detected with anti-CD3ζ antibodies (Fig. 3F). Cell lysis with Brij96, which preserves TCR nanoclusters, showed a notable reduction in large TCR complexes in CCR5^−/−^ compared to WT lymphoblasts (Fig. 3F). Two independent techniques thus support a CCR5 role in TCR nanoscopic organization in antigen-experienced CD4^+^ T cells.

To determine whether CCR5 controls TCR nanoclustering in *in vivo*-generated memory T cells, we analyzed TCR distribution in surface replicas of CD4^+^ memory T cells purified by negative selection from OVA/OVA-immunized WT and CCR5^−/−^ mice (Fig. S3). CD4^+^ memory cells from CCR5^−/−^ mice showed fewer, smaller TCR nanoclusters than those from WT counterparts (Fig. 3G), which indicates that CCR5 promotes formation of large TCR nanoclusters in endogenously generated CD4^+^ memory T cells.

### CCR5-induced TCR nanoclustering is independent of its co-stimulatory activity

Since CCR5 has co-stimulatory functions in CD4^+^ T cell priming (González-Martín et al., 2011, Molon et al., 2005), it is of interest to know whether defective TCR clustering in CCR5^−/−^ lymphoblasts is due to suboptimal primary activation of these cells. To address this question, we treated OT-II WT cells with the CCR5 antagonist TAK-779 at various intervals throughout culture, and analyzed TCR nanoclusters in IL-2-expanded T lymphoblasts. TAK-779 addition during the priming phase (blockade of CCR5 co-stimulatory function) decreased the percentage of large TCR nanoclusters compared to untreated controls (Fig. 4A). TAK-779 treatment did not alter TCR clustering in OT-II CCR5^−/−^ cells (Fig. S4), which indicates that the TAK-779 effect on OT-II cells is CCR5-specific.

**Figure 4.**
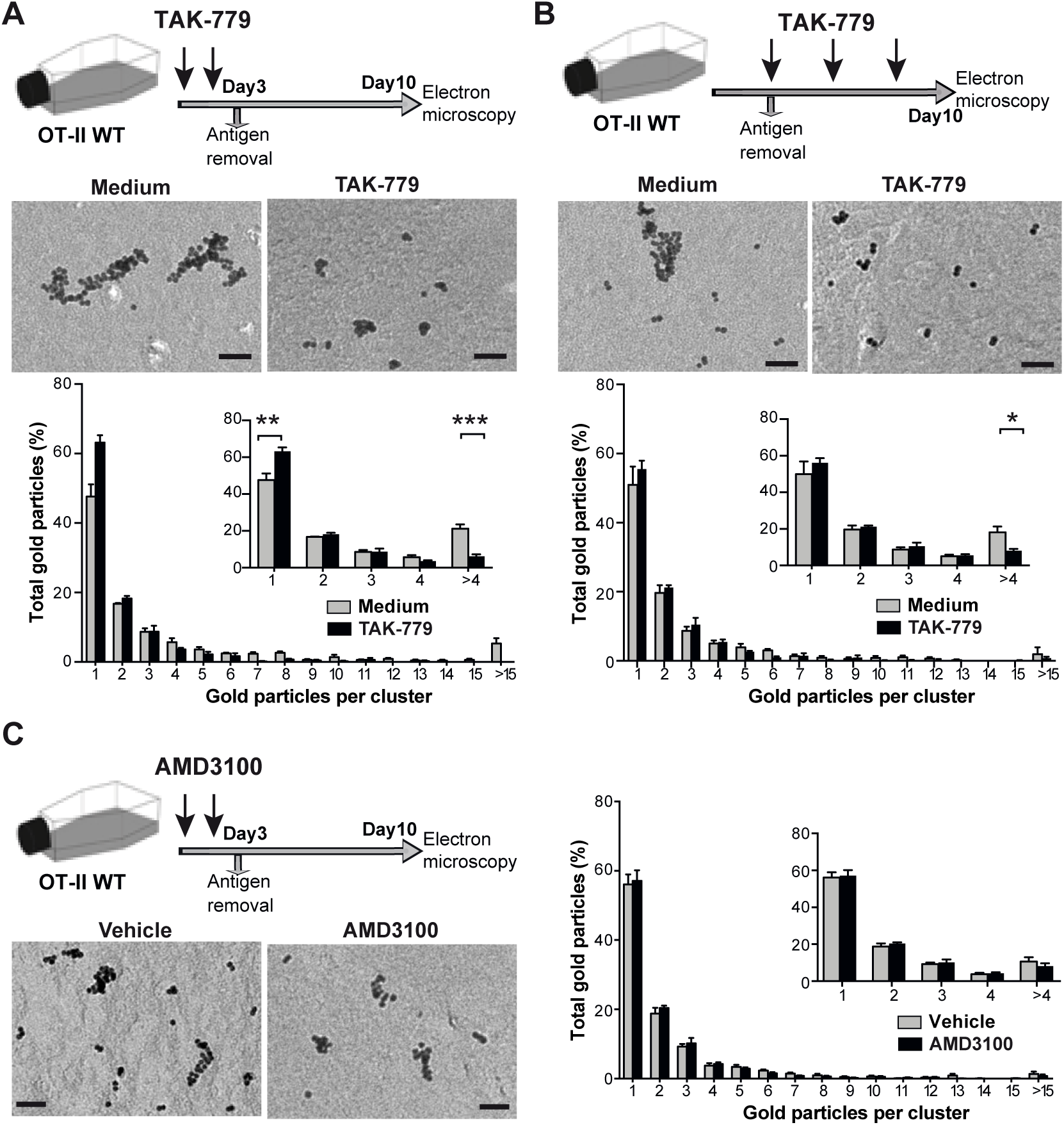
CCR5-induced TCR nanoclustering is specific and independent of its costimulatory activity. **A.** OT-II WT cells were activated with OVA_323-339_, alone or with TAK-779. After three days, antigen and TAK-779 were removed and lymphoblasts expanded in IL-2-containing medium. TCR nanoclustering was analyzed in anti-CD3ε-labeled surface replicas of day 10 lymphoblasts. Top, representative small field images showing gold particle distribution in the cell surface replicas of WT CD4^+^ T cells alone or with TAK-779 Bottom, quantification (mean ± SEM) of gold particles in clusters of the indicated size. Inset, distribution of gold particles between clusters of one, two, three, four or more than four particles in vehicle- (gray bars; *n* = 5 cells, 11266 particles) and TAK-779-treated cells (black; *n* = 6 cells, 5138 particles). **B.** OT-II WT cells were activated with OVA_323-339_, and TAK-779 was added at days 3, 5 and 7 after antigen removal. Analysis as above, untreated (gray bars; *n* = 5 cells, 6400 particles) and TAK-779-treated cells (black; *n* = 6 cells, 7153 particles). **C.** OT-II WT naïve cells were activated with antigen alone or with the CXCR4 inhibitor AMD3100. Top, representative images showing gold particle distribution in the cell surface replicas. Bottom, quantification (mean ± SEM) of gold particles in clusters of indicated size in the vehicle- (gray bars; *n* = 6 cells, 12339 particles) and AMD3100-treated cells (black; *n* = 7 cells, 17059 particles). Insets, distribution between clusters of one, two, three, four or more than four particles, and statistical analysis. * p <0.05, ** p <001, *** p <0.001, one-tailed unpaired Student’s *t*-test. Bar, 50 nm.

To avoid interference with the CCR5 co-stimulatory activity, we primed OT-II WT cells in the absence of the inhibitor, and added TAK-779 only during IL-2-driven expansion of the CD4^+^ lymphoblasts. In these conditions, TAK-779 also reduced the percentage of large TCR nanoclusters (Fig. 4B), which indicates that the CCR5 signals that control TCR organization are independent of those involved in its co-stimulatory function.

We next explored whether other chemokine receptors involved in T cell activation control TCR nanoclusters in CD4^+^ T cells. CXCR4 is a paradigmatic chemokine receptor that also provides costimulatory signals (Kumar et al., 2006, Smith et al., 2013). We primed OT-II WT cells in the presence of the CXCR4 antagonist AMD3100, and analyzed TCR nanoclusters in IL-2-expanded T lymphoblasts. Vehicle- and AMD3100-treated cells showed similar TCR nanocluster distribution (Fig. 4D), which implies that CXCR4 blockade does not interfere with TCR nanoclustering.

### CCR5 deficiency increases ceramide levels in CD4^+^ T cells

We analyzed CCR5 regulation of TCR nanoclustering in CD4^+^ T cells, and found no differences between OT-II WT and CCR5^−/−^ cells in TCR/CD3 chain mRNA levels or in cell surface expression of the TCRα chain (Fig. S5). These data suggest that the reduction in TCR clustering in CCR5^−/−^ cells is not due to decreased TCR expression.

TCR nanoclustering is dependent on plasma membrane Chol and SM (Molnar et al., 2012), two lipids also necessary for CCR5 signaling (Mañes et al., 2001). OT-II WT and CCR5^−/−^ lymphoblasts expressed comparable levels of total Chol and SM species (Fig. 5A, B). OT-II CCR5^−/−^ lymphoblasts nonetheless showed a significant increase in most ceramide (Cer) species and their dihydroCer (dhCer) precursors (Fig. 5C, D). These differences were not observed in naïve OT-II WT and CCR5^−/−^ cells (Fig. S6A), indicative that the Cer increase was specific to antigen-experienced cells. The increase in Cer species in CCR5^−/−^ lymphoblasts was not linked to enhanced apoptosis compared to WT cells (Fig. S6B).

**Figure 5.**
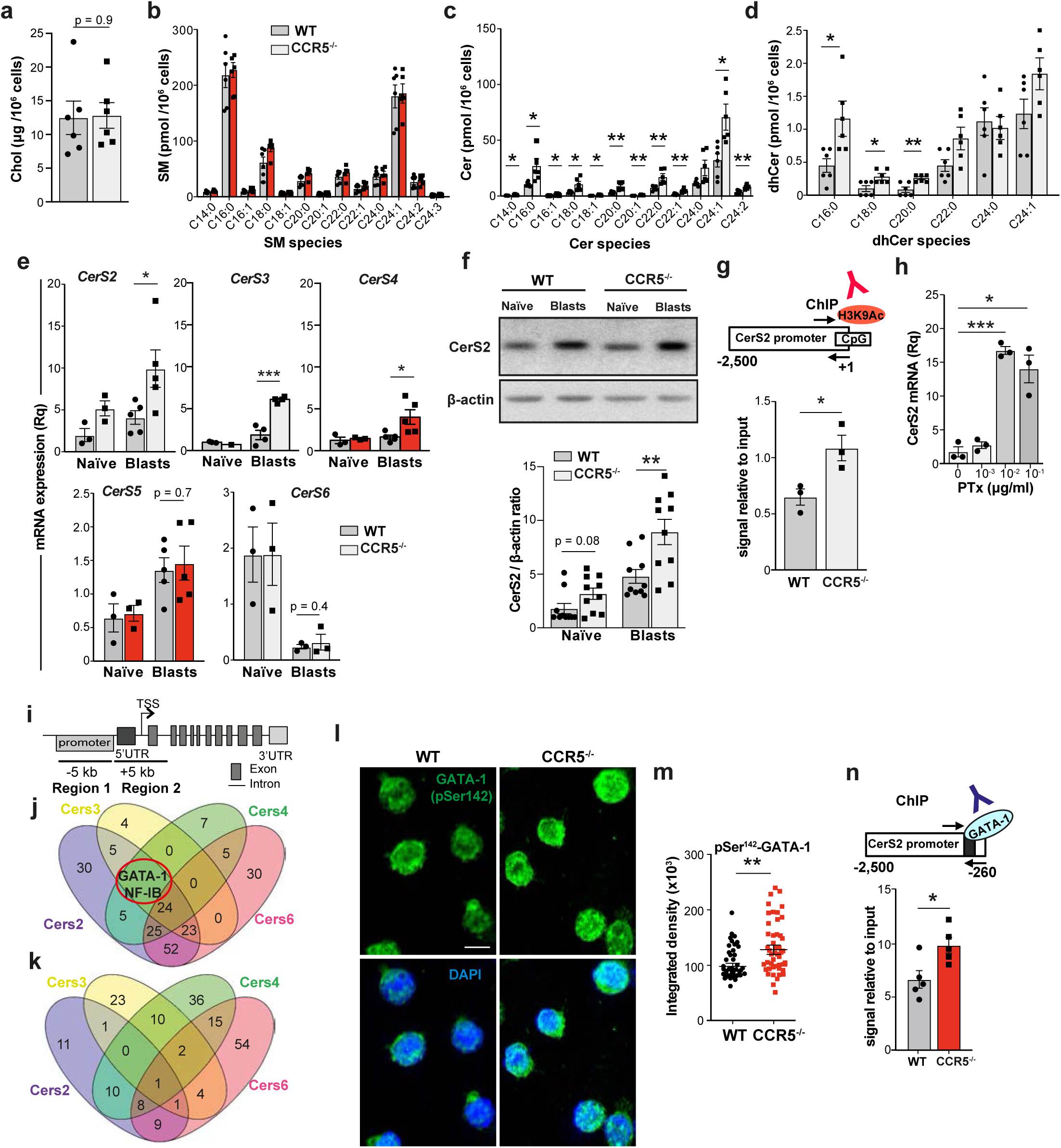
CCR5 deficiency increases Cer levels by upregulating specific CerS. **A.** Total Chol levels in WT and CCR5^−/−^ OT-II lymphoblasts (day 10, IL-2-expanded) as determined by a fluorometric assay. Data shown as mean ± SEM (*n* = 6). **B-D.** SM (*B*), Cer (*C*) and dhCer (*D*) levels in WT and CCR5^−/−^ OT-II 10-day lymphoblasts, as determined by UPLC-TOF MS. Values, after normalization with C17 standards and cell number in each sample, are the mean of two independent experiments (*n* = 6). **E.** RT-qPCR determination of CerS mRNA levels in naïve and IL-2-expanded WT and CCR5^−/−^ OT-II 10-day lymphoblasts. Data shown as mean ± SEM of triplicates from 3-5 independent experiments. **F.** Top, representative immunoblot showing CerS2 protein levels in naïve and WT and CCR5^−/−^ OT-II 10-day lymphoblasts. Bottom, densitometric quantification of blots as above (*n* = 10). **G.** ChIP analysis of the CerS2 promoter using an anti-H3K9Ac antibody. Top, scheme of the CerS2 promoter showing CpG islands and primers used for amplification. Bottom, relative ChIP of the CerS2 promoter in WT and CCR5^−/−^ OT-II 10-day lymphoblasts. Data shown as mean ± SEM of triplicates from 3 independent experiments. **H.** Relative CerS2 mRNA level in CD4 T cells treated with PTx. Each data point is the average of triplicates in an independent experiment. **I.** Scheme of a canonical CerS gene to illustrate the *in silico* strategy used to search for CerS-specific transcription factors. **J, K.** Venn diagrams showing the number of transcription factors with putative binding sites in the indicated CerS genes in regions 1 (*J*) and 2 (*K*). **L.** Representative immunofluorescence images showing pSer142-GATA-1 staining of OT-II WT and CCR5^−/−^ lymphoblasts. **M.** Quantification of nuclear staining of the cells plotted as integrated density fluorescence intensity in DAPI-stained area; *n* ≥50 cells/condition). **N.** Top, basic scheme of the CerS2 promoter, indicating the putative GATA-1 binding site (blue) and location of the primers used for amplification in ChIP assays. Bottom, relative anti-GATA-1 ChIP levels in OT-II WT and CCR5^−/−^ lymphoblasts. Data shown as mean ± SEM of triplicates (*n* = 5). * p <0.05, ** p <001, two-tailed unpaired Student’s *t*-test.

### CCR5 deficiency upregulates specific ceramide synthases in CD4^+^ T cells

Our analysis of the mRNA levels of key enzymes involved in Cer metabolism showed no differences in ceramidases (*ASAH1, ACER 2, ACER 3*) and sphingomyelinases (*SMPD1-4*) between OT-II WT and CCR5^−/−^ naïve cells or lymphoblasts (Fig. S7). mRNA levels of the ceramide synthases (CerS) CerS2, CerS3 and CerS4 were nonetheless upregulated in OT-II CCR5^−/−^ lymphoblasts (Fig. 5E); CerS5 and CerS6 were unaltered, and the nervous system-specific CerS1 isoenzyme was not detected. CerS2, CerS3 and CerS4 levels were comparable in naïve CD4^+^ WT or CCR5^−/−^ cells (Fig. 5E), which again associates the CCR5 transcriptional effect on these genes with activation.

We sought to validate the CerS isoforms upregulated by CCR5 deficiency at the protein level. In accordance with mRNA analyses, CerS2 protein levels were significantly higher in CCR5^−/−^ than in WT lymphoblasts (Fig. 5F); CerS3 and CerS4 were undetectable or only barely detectable by immunoblot. This is consistent with the fact that CerS2 has the highest expression level and the broadest substrate specificity in other cell types (Laviad et al., 2008). Chromatin immunoprecipitation (ChIP), followed by amplification of a region of the CerS2 promoter enriched in CpG islands, showed that binding of the transcriptional activation marker acetylated histone H3K9 (H3K9Ac) was higher in CCR5^−/−^ than in WT lymphoblasts (Fig. 5G). Moreover, blockade of CCR5 signaling with pertussis toxin (PTx; an inhibitor of the Gα_i_ subunit) also increased CerS2 mRNA expression (Fig. 5H).

To further study CCR5 transcriptional regulation of CerS, we scanned for transcription factors with putative binding sites in the CerS2, CerS3 and CerS4 promoters, which are transcriptionally upregulated in CCR5^−/−^ lymphoblasts, but not represented in the CerS6 promoter, which is not CCR5-regulated. We selected two regions; region 1 comprised −5 kb to the 5’UTR, and region 2 encompassed the 5’UTR to the first coding exon (Fig. 5I). This bioinformatic approach identified GATA-1 and NF-IB (nuclear factor-1B) as putative transcription factors involved in the differential expression of the CerS2 isoform (Fig. 5J, K).

We focused on GATA-1, since it is implicated in the differentiation of some CD4^+^ T cell subtypes (Fu et al., 2012, Sundrud et al., 2005). Immunofluorescence analyses showed increased nuclear levels of the phosphoSer142-GATA-1 form in OT-II CCR5^−/−^ compared to WT lymphoblasts (Fig. 5L, M), which correlated with enriched GATA-1 binding to the CerS2 promoter in CD4^+^ CCR5^−/−^ lymphoblasts (Fig. 5N). These results suggest that high Cer levels in CCR5^−/−^ lymphoblasts are a result of transcriptional induction of CerS2 through GATA-1.

### Ceramide levels control TCR nanoclustering

We used a synthetic biology approach to determine whether ceramide content affects TCR nanoclustering. Large unilamellar vesicles (LUV) were prepared at different molar ratios of PC, Chol, SM and Cer (Fig.6A), and then reconstituted with a streptavidin-binding-peptide-tagged TCR purified in its native state from murine M.mζ-SBP T cells (Swamy & Schamel, 2009). The proteoliposomes were analyzed by BN-PAGE after solubilization in 0.5% Brij96 to maintain TCR nanocluster integrity, or in 1% digitonin to disrupt TCR clusters. As anticipated (Molnar et al., 2012, Wang et al., 2016), TCR was monomeric in PC-containing LUV, whereas it formed nanoclusters when reconstituted in PC/Chol/SM liposomes (Fig. 6B, C). The inclusion of ceramides in these LUV (PC/Chol/SM/Cer liposomes) reduced TCR nanoclustering in a dose-dependent manner. This effect was not due to differential TCR reconstitution in Cer-containing LUV, since digitonin treatment rendered equivalent levels of monomeric TCR in each condition (Fig. 6B). These data suggest that Cer membrane content impairs TCR nanoclustering.

**Figure 6.**
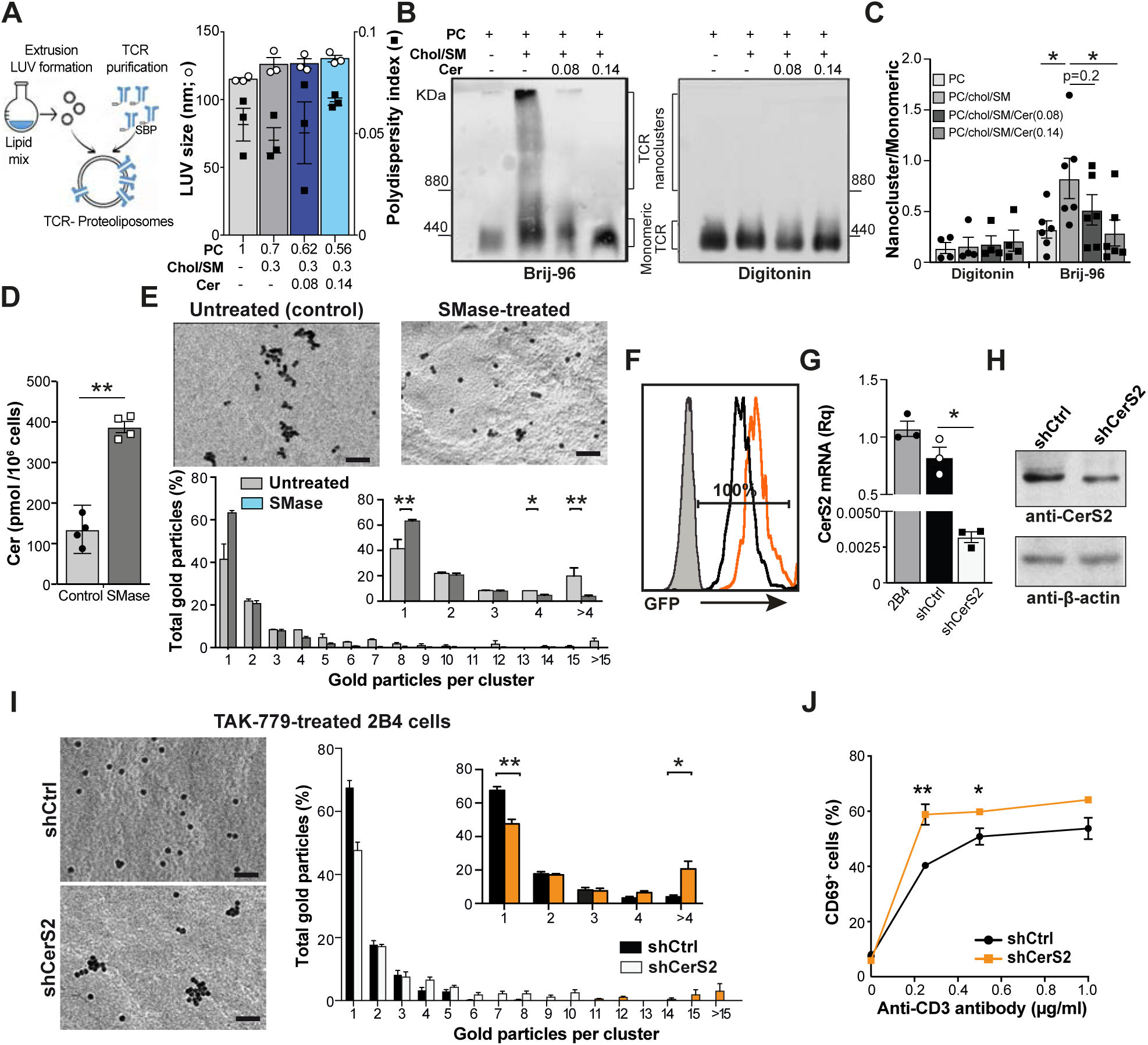
Cer levels determine the grade of TCR nanoclustering. **A.** Scheme of the strategy used to form TCR proteoliposomes, and size of LUV generated at the indicated lipid molar ratio. Polydispersity index values are shown as black squares for each condition (*n* = 3). **B.** Representative immunoblots comparing TCR nanocluster sizes *via* BN-PAGE and anti-CD3ζ immunoblotting in TCR proteoliposomes lysed in the presence of Brij-96 or digitonin. The marker protein is ferritin (f1, 440 and f2, 880 kDa forms). **C.** The ratio of the nanocluster and monomeric TCR in each lysis condition was quantified by densitometry from immunoblots as in *B*. Data shown as mean ± SEM of at least 4 independent experiments. **D.** Cer levels in OT-II 10-day lymphoblasts, untreated or treated with SMase (*n* = 2). **E.** Representative small field image showing gold particle distribution, and quantification (mean ± SEM) of gold particles in clusters of the indicated size in cell surface replicas from untreated (gray bars; *n* = 5 cells, 8126 particles) and SMase-treated (1h) OT-II lymphoblasts (black; *n* = 6 cells, 8457 particles) after CD3ε labeling, as determined by EM. The inset shows distribution between clusters of one, two, three, four or more than four particles, and statistical analysis. **F.** GFP expression in shCtrl- (black) and shCerS2 (orange)-transduced 2B4 cells after puromycin selection, as determined by FACS. **G.** Relative CerS2 mRNA levels in TAK-779-treated 2B4 cells as in *F*. Values were normalized to those obtained in untransduced TAK-779-treated 2B4 cells. Data shown as mean ± SEM (*n* = 3). **H.** Representative immunoblot with anti-CerS2 antibody to determine CerS2 protein levels in shCtrl and ShCerS2-transduced 2B4 cells as in *G*. Filters were rehybridized with β-actin as loading control. **I.** TCR nanoclustering of shCtrl- and shCerS2 transduced 2B4 cells in the presence of TAK-779. Representative small field images and quantification (mean ± SEM) of gold particles in clusters of indicated sizes in cell surface replicas shCtrl (gray bars; *n* = 6 cells, 12337 particles) and shCerS2 2B4 lymphoblasts (black; *n* = 7 cells, 13456 particles). Inset, distribution between clusters of indicated size, and statistical analysis. **J.** Percentage of CD69^+^ shCtrl (black) and shCerS2 (orange) 2B4 cells restimulated with plate-bound anti-CD3ε antibody in the presence of TAK-779. Data shown as mean ± SEM. * p <0.05, ** p <001, one-tailed (*I*) or two-tailed unpaired Student’s *t*-test. Bar, 50 nm.

To test whether this effect also occurs in live cells, we treated OT-II WT lymphoblasts with recombinant sphingomyelinase (SMase), which hydrolyzes SM to ceramide (Kitatani et al., 2008). SMase treatment of OT-II blasts increased Cer levels robustly (Fig. 6D), but did not compromise cell viability (Fig. S8). Analysis of membrane replicas from these cells showed that SMase treatment reduced the number of high valency TCR nanoclusters compared to controls (Fig. 6E), which indicates that high Cer levels hinder TCR nanoclustering in CD4^+^ T cells.

### CerS2 silencing restores TCR nanoclustering after CCR5 functional blockade

To correlate increased CerS2 expression with the impaired TCR nanoclustering in OT-II CCR5^−/−^ T cells, we attempted to silence CerS2 expression by lentiviral transduction of primary lymphoblasts with short-hairpin (sh) RNA for CerS2 or control (shCtrl). In the most successful experiments, we were only able to transduce ∼20% of the lymphoblasts, which did not lead to solid CerS2 mRNA silencing (Fig. S9A,B). Despite the low efficiency, antigenic restimulation tended to promote stronger responses in shCerS2-than in shCtrl-transduced cells (Fig. S9C,D). The low efficiency also precluded analysis of TCR nanoclusters in membrane replicas, as transduced cells could not be distinguished from non-transduced cells.

To overcome these difficulties, we used the 2B4 CD4^+^ T cell line. We verified that 2B4 cells expressed CCR5, and that TAK-779 treatment increased CerS2 levels and impaired TCR nanoclustering (Fig. S10). The data suggest that CCR5 effects on TCR nanoclustering and CerS2 induction are not exclusive to the OT-II system, and that TAK-779-treated 2B4 cells mimic the functional findings in OT-II CCR5^−/−^ lymphoblasts. 2B4 cells were transduced efficiently by lentiviruses and, after three days of antibiotic selection, 100% of the cells expressed the shRNA; this led to strong silencing of CerS2 mRNA and protein in shCerS2-compared to shCtrl-transduced cells (Fig. 6F-H). Analysis of TCR organization showed recovery of large TCR nanoclustering in TAK-779-treated, shCerS2-transduced cells compared to controls (Fig. 6I); after restimulation with plate-bound anti-CD3ε antibody in the presence of TAK-779, CD69 upregulation was higher in CerS2-deficient than in shCtrl-cells (Fig. 6J).

### CCR5 modulates TCR nanoclustering in human CD4^+^ T cells

Finally, we tested whether CCR5 deficiency also impairs TCR organization in human CD4^+^ T cells. Approximately 1% of the Spanish population bears the *ccr5*Δ32 polymorphism in homozygosity (Mañes et al., 2003). Purified CD4^+^ T cells from healthy WT or *ccr5*Δ32 homozygous donors were activated with anti-CD3 and -CD28 antibodies for three days, and maintained for five additional days with IL-2. We found that *ccr5*Δ32 lymphoblasts had a lower percentage of large TCR nanoclusters than WT cells (Fig. 7A); concomitantly, the fraction of monomeric TCR was increased in the former. Sphingolipid analysis of these CD4^+^ lymphoblasts showed an increase in saturated 24-carbon Cer (C24:0) and its precursor (dhCer C24:0) in cells derived from *ccr5*Δ32 donors, whereas SM levels were comparable between both genotypes (Fig. 7B). This increase in Cer levels was associated with upregulation of CerS2 mRNA in *ccr5*Δ32 lymphoblasts compared to WT controls (Fig. 7C); expression of other enzymes involved in Cer metabolism was unchanged in both genotypes (Fig. S11). As found in mouse CCR5^−/−^ lymphoblasts, antigen-experienced human CD4^+^ T cells from *ccr5*Δ32 homozygotes also show defective TCR nanoclustering associated with increased Cer levels and upregulated CerS2. Moreover, they indicate that these CCR5 effects are not restricted to specific T cell clones, but can be observed in a polyclonal T cell repertoire.

**Figure 7.**
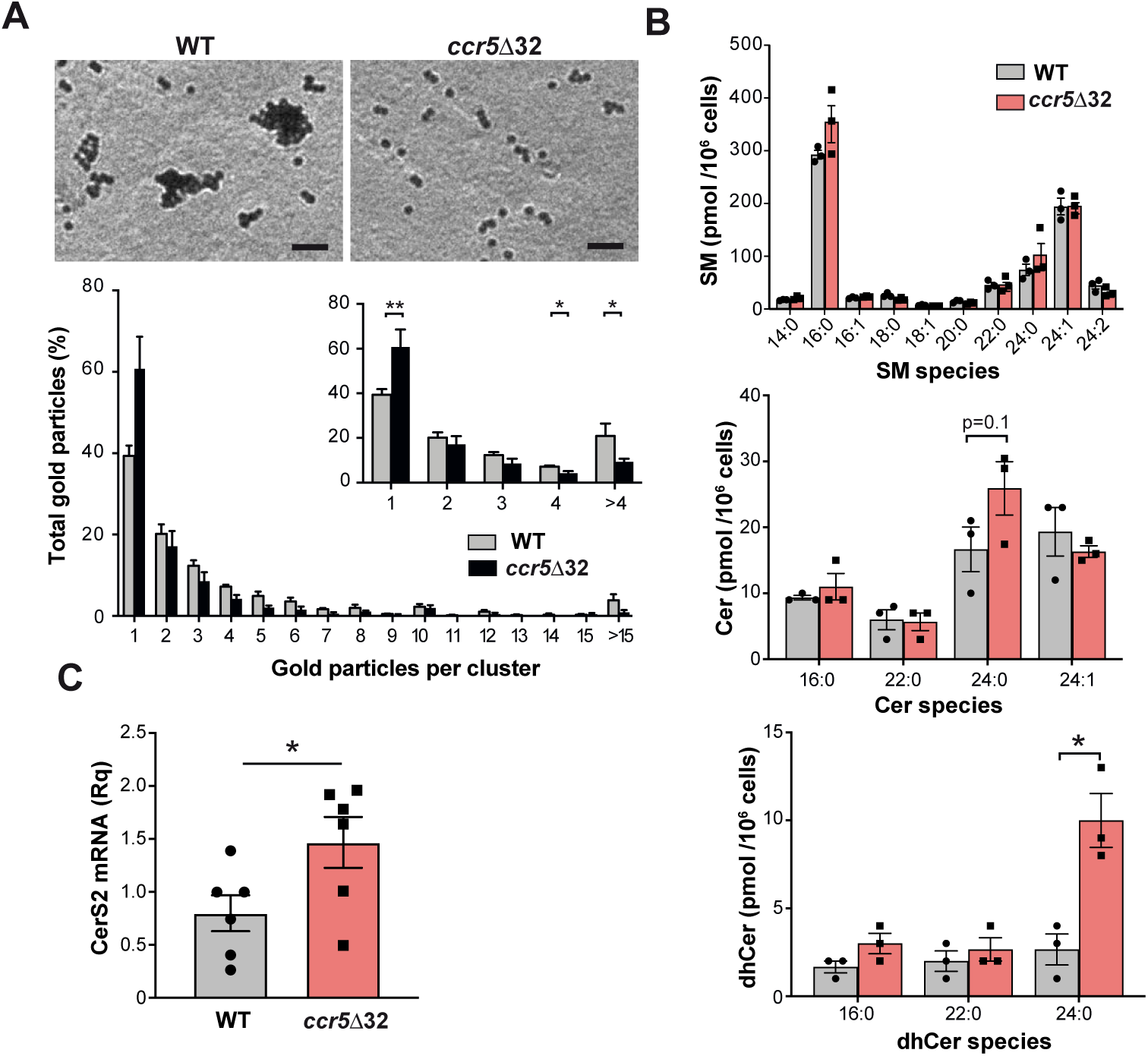
CCR5 controls TCR nanoclustering and Cer metabolism in human CD4^+^ cells. **A.** Analysis of TCR nanoclustering in lymphoblasts from healthy WT and *ccr5*Δ32 homozygous donors. Top, representative small field image showing gold particle distribution in cell surface replicas of anti-CD3ε-labeled cells; bottom, quantification (mean ± SEM) of gold particles in clusters of the indicated size in the WT (gray bars; *n* = 5 cells, 17689 particles) and Δ32/Δ32 cells (black; *n* = 4 cells, 16938 particles). Replicas were generated from 3 donors of each genotype. Insets show the distribution between clusters of one, two, three, four or more than four particles, and statistical analysis. **B.** Normalized SM, Cer and dhCer levels in lymphoblasts obtained as in *A*. Data shown as mean ± SEM from three donors of each genotype. **C.** Relative CerS2 mRNA levels in day 8 WT and *ccr5*Δ32 lymphoblasts. Data shown as mean ± SEM of triplicates from 3 donors in two independent experiments. * p <0.05, ** p <001, two-tailed unpaired Student’s *t*-test. Bar, 50 nm.

## DISCUSSION

Here we show that CCR5 signaling is largely dispensable for memory CD4^+^ T cell differentiation, but provides specific signals that improve the functional fitness of memory cells after antigen re-encounter. The CCR5 signals optimize TCR nanoclustering and antigen sensitivity by triggering a CD4^+^ T cell-specific transcription program that regulates Cer metabolism. This CCR5 program operates in murine and human CD4^+^ T cells, which suggests physio-pathological relevance.

A central observation of our study is that CCR5 expression enhances the degree of TCR nanoclustering in resting antigen-experienced T cells, both *in vitro* and *in vivo*. The presence of TCR nanoclusters in resting T cells was shown by BN-PAGE, EM, and super-resolution microscopy (Hu et al., 2016, Jung et al., 2016, Pageon et al., 2016, Schamel et al., 2005). We demonstrate here differential TCR nanoclustering in WT and CCR5-deficient cells using two complementary approaches (EM and BN-PAGE), based on different conceptual principles. In EM, TCR nanoclusters were defined as gold particle aggregates at less than 10 nm distance from one another. Previous analyses showed that this criterion permits identification of TCR nanoclusters formed by TCR-TCR interactions (Kumar et al., 2011); these tightly-associated TCR nanoclusters would allow inter-TCR cooperativity for pMHC binding (Martín-Blanco et al., 2018). We therefore intentionally considered TCR not to be in the same nanocluster if the gap between them was >10 nm; this excludes considering more loosely associated TCR as nanoclusters, but the strict definition allowed association of TCR nanoclusters to a T cell biological function.

It is also important to clarify that gold particle counts do not necessarily correspond to the number of TCR molecules in a nanocluster. BN-PAGE defines neither the exact size nor the abundance of TCR nanoclusters. Direct comparison of CCR5^−/−^ with WT cells using both methods nonetheless allowed us to detect relative differences and determine the promoter effect of CCR5 in TCR nanoclustering. Application of Monte Carlo simulations further indicated that the nanoclusters observed in EM are not the result of random proximity of gold particles. The estimated clustering parameter (*b*) for randomly distributed particles was virtually zero.

Since CCR5 provides positive signals during activation of naïve CD4^+^ T cells (González-Martín et al., 2011, Molon et al., 2005, Nesbeth et al., 2010), we attempted to clarify whether TCR nanoclustering impairment in CCR5^−/−^ cells is solely an effect of this defective priming. This is unlikely, since TCR clustering was inhibited when CCR5 was blocked during expansion of fully-activated WT lymphoblasts. This effect on TCR nanoclustering during lymphoblast expansion was modest compared to that observed in the priming phase, but is probably the result of insufficient CCR5 inhibition during lymphoblast expansion. CCR5 is not only upregulated shortly after activation, but is maintained in memory CD4^+^ T cells, which are highly susceptible to infection by R5-tropic HIV-1 strains (Nie et al., 2009). Our results showed increased CCR5 mRNA expression during lymphoblast expansion. These lymphoblasts also expressed CCR5 ligands, suggesting autocrine CCR5 stimulation during this phase. We thus propose that TCR nanoclustering is regulated by CCR5 signals transduced during lymphoblast differentiation rather than during priming.

Another feature that distinguishes CCR5 effects on priming and on TCR nanoclustering is the role of CXCR4 in these events. During priming, CCR5 and CXCR4 are recruited to and accumulate as heterodimeric complexes at the IS of CD4^+^ T cells; AMD3100 (a CXCR4 antagonist) prevented not only CXCR4 but also CCR5 accumulation (Contento et al., 2008), which indicates necessary cooperation between CCR5 and CXCR4 for full T cell activation. In contrast, AMD3100 did not affect TCR nanoclusters in CCR5-expessing cells, which suggests that CXCR4/CCR5 heterodimer signaling is not essential for TCR nanoclustering in lymphoblasts. CCR5 homo- and heterodimers are thought to associate differently to Gα subunits; homodimers signal through the PTx-sensitive Gα_i_, whereas heterodimers generate PTx-resistant responses (Mellado et al., 2001). CCR5 costimulatory signals in the IS are PTx-resistant (Molon et al., 2005), consistent with CCR5/CXCR4 heterodimerization during priming. PTx potentiates transactivation of the CerS2 promoter (Fig. 5H), however, which suggests involvement of the CCR5-induced Gα_i_ pathway in TCR nanoclustering. CCR5 thus appears to trigger distinct signaling pathways for co-stimulation and TCR nanoclustering in CD4^+^ T cells.

Cholesterol and SM are two lipids essential for CCR5 signaling and TCR nanoclustering (Mañes et al., 2001, Molnar et al., 2012). In resting T cells, these receptors nonetheless partition in different membrane phases, liquid-ordered (l_o_) for CCR5 (Molon et al., 2005) and liquid-disordered (l_d_) for TCR (Beck-Garcia et al., 2015). This differential phase segregation argues against direct CCR5/TCR interaction as a mechanism that influences TCR nanoclustering. Our results suggest instead that increased levels of long-chain Cer species cause defective TCR nanoclustering in CCR5^−/−^ lymphoblasts. Indeed, elevation of Cer levels in TCR-reconstituted proteoliposomes and in live cells by SM hydrolysis impaired nanoscopic TCR organization. Although Cer levels were higher in lymphoblasts than in naïve cells, which supports a role for Cer in T cell activation (Sofi et al., 2017), CCR5 deficiency further increased Cer levels specifically in lymphoblasts. The Cer increase in CCR5^−/−^ lymphoblasts did not cause spontaneous apoptosis (Fig. S6B), which coincides with the non-apoptotic and preventive effects of long-chain Cer in this process (Stiban & Perera, 2015). We propose that in activated CD4^+^ T cells, autocrine CCR5 activation downgrades the nuclear entry and binding of activated GATA-1 to specific CerS promoters, such as that of CerS2; this restrict Cer biosynthesis and provide a membrane environment permissive for TCR nanoclustering (Fig. 8). Given that WT and CCR5^−/−^ lymphoblasts have similar SM levels, our results pinpoint *de novo* Cer biosynthesis as an important metabolic checkpoint for CD4^+^ T cell memory function.

**Figure 8.**
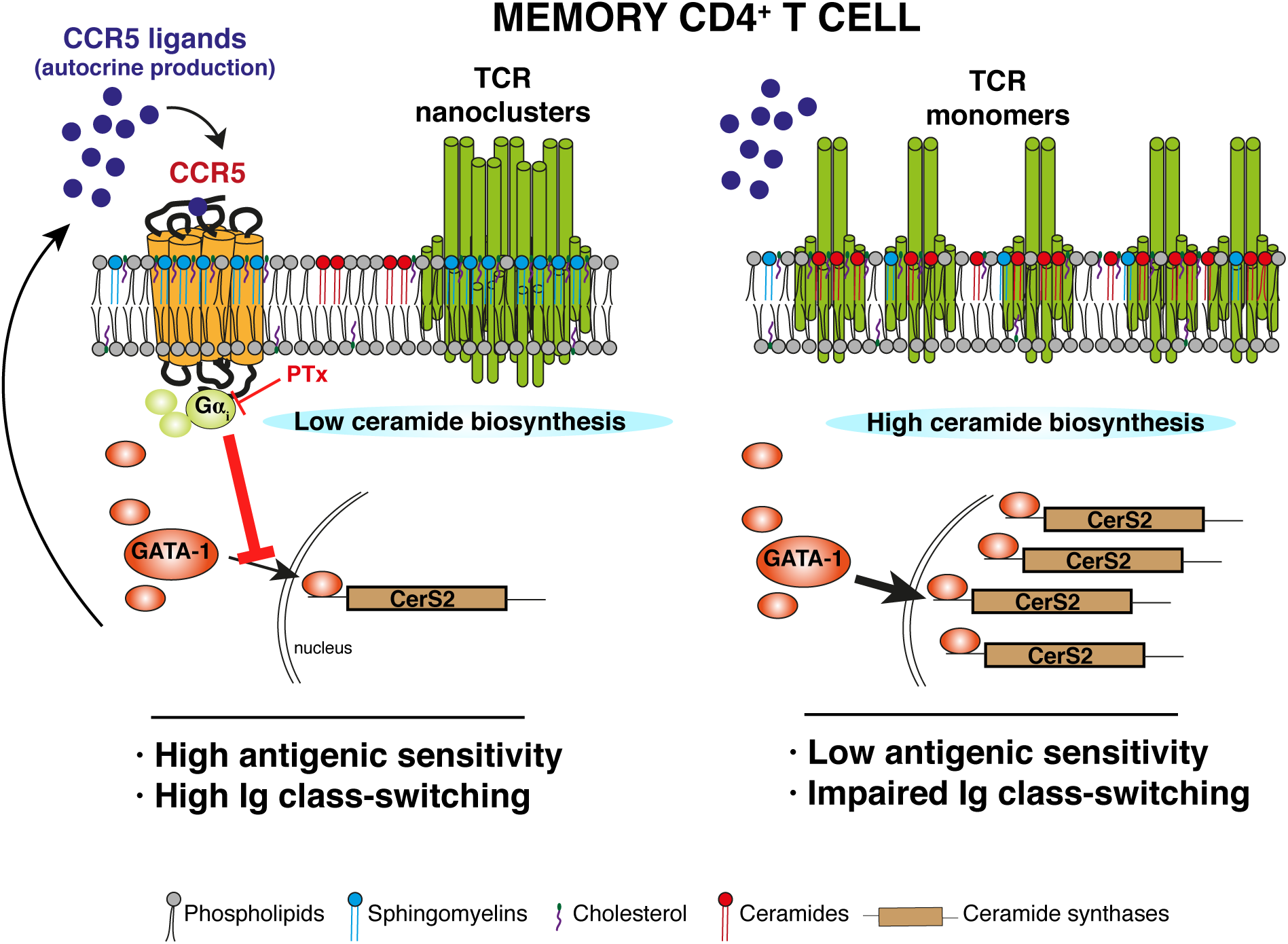
Scheme showing the CCR5 effect on TCR nanoclustering in antigen-experienced CD4^+^ T cells. Autocrine activation of CCR5 provides antigen-independent signals that regulate TCR nanoclustering in antigen-experienced CD4^+^ T cells, by limiting GATA-1-induced transcription of CerS2; this maintains control of the *de novo* Cer biosynthetic pathway and restrains Cer levels (left panel). CCR5 signals thus endow CD4^+^ lymphoblasts with a plasma membrane lipid environment that supports maximal TCR nanoclustering, which increases antigenic sensitivity and improves T:B cell cooperation for humoral responses after antigen re-encounter. In antigen-experienced CD4^+^ T cells that lack CCR5, such as those from *ccr5*Δ32 homozygous individuals (right panel), the restriction of GATA-1-induced CerS2 transcription is lost. This leads to aberrant activation of Cer biosynthesis and, through an increase in Cer levels, to generation of a restrictive lipid environment for TCR nanoclustering. As a result, antigenic sensitivity and humoral responses are impaired in CCR5-deficient, antigen-experienced CD4^+^ T cells.

Several settings can be hypothesized that explain Cer effects on TCR nanoclustering. In model membranes, Cer have strong segregation capacity, which might affect their lateral organization (Goñi & Alonso, 2009). When the Chol concentration is saturating, however, as is the case of cell membranes and our proteoliposomes, Cer-enriched domains are not formed, due to the ability of Chol and Cer to displace one another (Castro et al., 2009). This argues against the idea that Cer impairs TCR clustering by promoting a general reduction in membrane lateral diffusion. Our mathematical model also predicted that cluster distribution is independent of TCR diffusivity. Interaction of the TCRβ subunit with Chol/SM complexes is critical for TCR nanoclustering (Molnar et al., 2012, Wang et al., 2016). High levels of Cer or their precursors (dhCer) could transform Chol/SM into Chol/SM/Cer domains with specific physicochemical properties, which might hinder TCR nanocluster formation physically or thermodynamically. For instance, high dhSM levels inhibit CCR5-mediated HIV-1 infection by rigidifying CCR5-containing l_o_ domains (Vieira et al., 2010). Chol/SM interaction depends on the hydrogen bond generated by the amide group of the SM molecule and the 3-hydroxyl group of Chol (Ramstedt & Slotte, 2002), but the SM amide group can also form hydrogen bonds with the Cer hydroxyl group (Garcia-Arribas et al., 2016). It is thus possible that, rather than forming SM/Chol/Cer domains, small increases in Cer levels would increase the mutual displacement of three lipids. This could lead to replacement of SM/Chol by SM/Cer complexes with gel-like biophysical properties (Sot et al., 2008).

Our results indicate that the CCR5 effect on TCR clustering is biologically meaningful. In a first model, we show that the responses of CCR5-deficient memory CD4^+^ T cells generated by vaccination were impaired after *ex vivo* stimulation. In a second model that involves T:B cell cooperation, we show that CCR5 deficiency impaired class switching of high affinity antibodies after re-exposure to a T cell-dependent antigen. Affinity maturation and class switching depend on recruitment of T_fh_ cells to GC (Vinuesa et al., 2016). This T_fh_ cell confinement is a result of CXCR5 expression and the downregulation of other homing receptors, including CCR5 (Crotty, 2011), which could explain the lack of difference in class switching between OVA/KLH-immunized WT and CCR5^−/−^ mice. There were also no differences between WT and CCR5^−/−^ effector T_fh_ cells after the first OVA immunization. Once GC resolve, however, some T_fh_ cells are reported to enter the circulation as T_fh_ central memory-like cells (Vinuesa et al., 2016). These circulating, antigen-experienced T_fh_ cells express CCR5 and are very susceptible to HIV-1 infection (Xu et al., 2017). We found that the frequency of large TCR nanoclusters increased in memory T cells from WT compared to CCR5^−/−^ OVA/OVA-immunized mice, which suggests increased antigenic sensitivity.

We hypothesize that following re-exposure to antigen, CCR5-expressing memory pre-T_fh_ cells will have a more efficient response than CCR5-deficient cells, which would support robust antibody responses after their differentiation to GC-T_fh_ cells. In humans, functional CCR5 deficiency does not cause strong immune suppression, but *ccr5*Δ32 homozygosity was associated with four times more fatal infections than average during the 2009-2011 influenza season in Spain (Falcon et al., 2015) and fatal infections by the West Nile Virus in the US (Lim & Murphy, 2011). Our results provide a conceptual framework on which to base clinical trials to evaluate CD4^+^ T cell memory responses in CCR5-deficient humans, and suggest caution regarding the risks associated to genetic ablation of CCR5 as a preventive strategy to block HIV-1 infection.

## MATERIAL AND METHODS

Resource Identification Portal accession numbers for antibodies, cell lines, animals and other reagents used in the study are provided as supplementary material.

### Antibodies and reagents

Antibodies used for characterization of mouse cells by flow cytometry were anti-Vα_2_TCR-PE (B20.1), -CD25-PE (PC61), -CD45.2-FITC (104), -CD62L-FITC/APC (MEL-14), -CD69-PeCy7 (H1.2F3) and biotinylated anti-CXCR5 (2G8) from BD-Biosciences; anti human biotinylated CD3 (OKT3), anti-CD4-PeCy7/efluor450/PacificBlue (RM4.5), -IFNγ-APC (XMG1.2), and -PD1-efluor780 (J43) from eBioscience, and anti-CD44-PeCy5/APC (IM7) from BioLegend. Biotinylated and purified anti-CD3ε (145-2C11; BD Biosciences) were used for electron microscopy and T cell activation, respectively. Anti-mouse CerS-2 (1A6; Novus Biologicals), -CD3ζ (449, purified from hybridoma), and -β-actin (AC-15; Sigma-Aldrich) were used for immunoblot. Anti-mouse GATA-1 (D52H6; Cell Signaling) and anti-mouse phospho-GATA1^pSer142^ (Thermo Fisher Scientific) were used for immunofluorescence. Anti-GATA1 (ab11852, Abcam), -histone H3Lys9 (CS200583) and purified IgG rabbit (PP64B; EMD Millipore) were used for ChIP.

The OVA_323–339_ peptide was synthesized at the CNB Proteomics facility. TAK-779, AMD-3100, poly-L-lysine, pertussis toxin, Cer (bovine spinal cord), and sphingomyelinase (*Bacillus cereus*) were from Sigma Aldrich. mCCL4, mIL-2, and mIL-15 were from Peprotech; NIP-KLH, NIP-OVA, NIP(7)-BSA, and NIP(41)-BSA from Biosearch Technology. Soybean phosphatidylcholine Chol, egg SM, C12 Cer (d18:1/12:0), C16 Cer (d18:1/16:0), C18 Cer (d18:1/18:0), C24 Cer (d18:1/24:0), C24:1 Cer (d18:1/24:1(15Z)), C16 dhCer (d18:0/16:0), C18 dhCer (d18:0/18:0), C24 dhCer (d18:0/24:0), C24:1 dhCer (d18:0/24:1(15Z)), C12:0 SM (d18:1/12:0), C16:0 SM (d18:1/16:0), C18:0 SM (d18:1/18:0), C24:0 SM, C24:1 SM, and the Cer mix from bovine spinal cord were from Avanti Polar Lipids. Lentiviral pGIPZ containing shRNAs for murine Cers 2 (V3LMM_454307, V3LMM_454309, and V3LMM_454311 clones), and the mismatched control were from Dharmacon.

### Mice and cell lines

C57BL/6J wild-type (WT) and CCR5^−/−^ mice were from The Jackson Laboratory. TCR transgenic OT-II CCR5^−/−^ mice, recognizing OVA_323-339_ (ISQAVHAAHAEINEAGR; I-Ab MHC class II molecule) have been described (González-Martín et al., 2011). B6-SJL (Ptprca Pepcb/BoyJ) mice bearing the pan-leukocyte marker allele CD45.1 were used for adoptive transfer experiments. CD3ε-deficient mice (DeJarnette, Sommers et al., 1998) were used as a source of antigen-presenting cells for restimulation assays. Mice were maintained in SPF conditions in the CNB and CBM animal facilities, in accordance with national and EU guidelines. All animal procedures were approved by the CNB and the Comunidad de Madrid ethical committees (PROEX 277/14; PROEX 090/19). Human embryonic kidney HEK-293T cells and the murine 2B4 hybridoma and its derivative M.mζ-SBP (which expresses a SBP-tagged form of CD3ζ) (Swamy & Schamel, 2009) were cultured in standard conditions.

### Isolation and culture of mouse and human primary T cells

Spleen and lymph nodes from 6- to 12-week-old OT-II WT and CCR5^−/−^ mice were isolated and cell suspensions obtained using 40 µM pore filters. Erythrocytes were lysed with AKT lysis buffer (0.15 M NH_4_Cl, 10 mM KHCO_3_, 0.1 mM EDTA), and cells activated with the appropriate OVA peptide for three days. Antigen was removed and cells were cultured with IL-2 (5 ng/mL) or IL-15 (20 ng/mL). For some experiments, naïve OT-I and OT-II cells were obtained by negative selection using the Dynabeads Untouched Mouse CD4 cell kit (ThermoFisher). Flow cytometry indicated >85% enrichment in all cases. Memory CD4^+^ T cells, generated *in vivo* after NIP-OVA or NIP-KLH immunization (see below), were isolated by negative selection with the Mouse Memory T cell CD4^+^/CD62L^−^/CD44^hi^ Column Kit (R&D Systems).

Blood samples from *ccr5*Δ32 homozygous and WT healthy donors were from the Fundació ACE (Barcelona, Spain), obtained with informed consent of the donors. No personal data were registered, and all procedures using these samples were in accordance with the standards approved by the Ethics Committee of the Hospital Clinic Barcelona (HCB/2014/0494 and HCB/2016/0659). Human peripheral blood mononuclear cells (PBMC) were isolated from Vacutainer Cell Preparation Tubes by separation on a Ficoll gradient. CD4^+^ T cells were obtained by negative selection using the EasySep Human CD4^+^ Enrichment kit (Stem Cell Technologies), and stimulated with anti-CD3-coated magnetic beads (Dynabeads M-450 tosyl-activated, Thermo Fisher) for 3 days. Beads were removed with a magnet and cells were incubated with IL-2 (5 ng/ml) to generate lymphoblasts. The *ccr5*Δ32 polymorphism (rs333) was genotyped by PCR (AriaMx Real-time; Agilent Technologies) as described (Mañes et al., 2003).

### Flow cytometry

For cell surface markers, cell suspensions were incubated (20 min, 4°C) with the indicated fluorochrome-labeled or biotinylated monoclonal primary antibodies in phosphate-buffered saline with 1% BSA and 0.02% NaN_3_ (PBS staining buffer). For intracellular labeling, cells were fixed and permeabilized with IntraPrep (Beckman Coulter), followed by intracellular staining with indicated antibodies. Cells were analyzed on Cytomics FC500 or Gallios cytometers (both from Beckman-Coulter) and data analyzed using FlowJo software.

### Immunization and adoptive transfer

Spleen and lymph node cell suspensions from OT-II WT or CCR5^−/−^ cells were adoptively transferred (5 × 10^6^ cells/ mouse) into CD45.1 mice. The following day, recipient mice were infected intravenously with rVACV-OVA virus (2 × 10^6^ pfu). Mice were sacrificed 35 days later, and splenocyte suspensions obtained as described above.

C57BL/6J or CCR5^−/−^ mice were immunized (i.p.) with NIP-OVA (200 µg) in alum (100 µl) diluted 1:1 in PBS. At 7 days post-immunization, spleens from three mice of each genotype were harvested and analyzed by flow cytometry to detect T_fh_ cells. On day 30, half of the mice in each group were re-immunized with NIP-OVA/alum (as above); the other half received NIP-KLH (200 µg)/alum. Mice were sacrificed 15 days later, and serum anti-NIP antibodies determined by ELISA. Plate-bound NP(7)-BSA and NP(41)-BSA (5 µg/ml) were used to measure high- and low-affinity Ig, respectively. Sera from NIP-OVA- and NIP-KLH-immunized mice were diluted 1:175 and, after several washing steps, anti-NIP antibody binding was developed with the SBA Clonotyping System-HRP (Southern Biotech). Absorbance at 405 nm was determined in a FilterMax F5 microplate reader (Molecular Devices). Memory cells from NIP-OVA- and NIP-KLH-immunized mice were purified as indicated and processed for EM.

### Immunogold labeling, replica preparation, and EM analysis

Immunogold-labeled cell surface replicas were obtained as described (Kumar et al., 2011). Briefly, T cells were fixed in 1% paraformaldehyde (PFA) and labeled with anti-mouse CD3 mAb (145-2C11) or anti-human CD3 mAb OKT3, followed by 10 nm gold-conjugated protein A (Sigma-Aldrich). Labeled cells were adhered to poly-L-lysine-coated mica strips and fixed with 0.1% glutaraldehyde. Samples were covered with another mica strip, frozen in liquid ethane (KF-80, Leica), and stored in liquid nitrogen. Cell replicas were prepared with a Balzers 400T freeze fracture (FF) unit, mounted on copper grids, and analyzed on a JEM1010 electron microscope (Jeol, Japan) operating at 80 kV. Images were taken with a CCD camera (Bioscan, Gatan, Pleasanton, CA) and processed with TVIPS software (TVIPS, Gauting, DE). Gold particles were counted on the computer. When distance between gold particles was smaller than their diameter (10 nm), they were considered part of the same cluster.

### BN-PAGE analysis of TCR clustering

Membrane fractions from OT-II WT and CCR5^−/−^ cells (20 × 10^6^) were prepared with a Dounce homogenizer, followed by incubation in hypotonic buffer (10 mM HEPES pH 7.4, 42 mM KCl, 5 mM MgCl_2_, protease inhibitors). Membranes were recovered by ultracentrifugation (100,000 xg, 45 min, 4°C) and lysed in 150 µl BN lysis buffer (500 mM Bis-Tris 40 mM pH 7.0, 1 mM ε-aminocaproic acid, 40 mM NaCl, 20% glycerol, 4 mM EDTA and 0.5% Brij96 or 1% digitonin) with protease inhibitors. BN-PAGE gradient gels (4–8%) were prepared and used as described (Swamy & Schamel, 2009), using ferritin 24-mer and 48-mer (f1, 440 kDa; f2, 880 kDa) as protein markers. Proteins were transferred to PVDF membranes and probed with anti-CD3ζ antibody.

### Restimulation assays

Splenocytes from CD3ε^−/−^ mice were irradiated (15 Gy), seeded (0.6 × 10^5^ cells/well) and loaded (2 h, 37°C) with different concentrations of OVA_323–339_ peptide. After centrifugation (300 xg, 5 min), isolated lymphoblasts (0.75 × 10^5^ cells/well) were co-cultured for 48 h. Supernatants were collected to measure IL-2 by ELISA (ELISA MAX Deluxe, BioLegend) and proliferation assessed by methyl-^3^[H]-thymidine (1 µCi/well) incorporation into DNA, in a 1450 Microbeta liquid scintillation counter (PerkinElmer).

### TCR purification

The TCR fused to streptavidin binding peptide was purified from M.mζ-SBP cells. Briefly, 100 × 10^6^ cells were lysed in lysis buffer (20 mM Bis-Tris pH 7, 500 mM ζ-aminocaproic acid, 20 mM NaCl, 2 mM EDTA, 10% glycerol, 1% digitonin). After incubation of the lysate with streptavidin-conjugated agarose (overnight, 4°C), the TCR was eluted by incubating samples with 2 mM biotin in lysis buffer (30 min, 4°C).

### Preparation of large unilamellar vesicles (LUV) and TCR reconstitution

LUV with a custom lipid composition were prepared by the thin film method (Molnar et al., 2012), followed by extrusion through polycarbonate membranes with a pore size of 200 nm (21 times) and 80 nm (51 times). The diameter of the resulting LUV was determined by dynamic light scattering (Zetamaster S, Malvern Instruments). The LUV preparation (2 mM) was mixed with purified TCR (0.1 µg) in 100 µl saline-phosphate buffer with 0.02% Triton X-100, and 40 µl 0.01% Triton X-100 was added. Samples were agitated (30 min, 4°C), and detergent removed by adsorption to polystyrene BioBeads SM-2 (3 mg; Bio-Rad; overnight, 4°C). Proteoliposomes were collected by ultracentrifugation (180,000 xg, 4 h, 4°C), lysed, and analyzed by BN-PAGE as above.

### Sphingolipid and Chol quantification

Total Chol level was measured with the Amplex Red cholesterol assay kit (Invitrogen) after lysis (50 mM Tris-HCl pH 8; 150 mM NaCl, 1% NP-40). Prior to sphingolipid quantification, we prepared calibration curves with mixtures of C12Cer, C16Cer, C18Cer, C24Cer, C24:1 Cer, C16dhCer, C18dhCer, C24dhCer, C24:1dhCer, C12SM, C16SM, C18SM, C24SM, and C24:1SM. For sphingolipid determination, cell pellets (1 × 10^6^) were mixed with internal standards (N-dodecanoyl-sphingosine, N-dodecanoylglucosylsphingosine, N-dodecanoylsphingosylphosphorylcholine, C17-sphinganine, and C17-sphinganine-1 phosphate; 0.2 nmol each; Avanti Polar Lipids) in a methanol:chloroform solution. Sphingolipids were extracted as described (Merrill, Sullards et al., 2005), solubilized in methanol, and analyzed by ultra-performance liquid chromatography (UPLC; Waters, Milford, MA) connected to a time-of-flight detector (TOF; LCT Premier XE) controlled by Waters/Micromass MassLynx software. Lipid species were identified based on accurate mass measurement with an error <5 ppm and their LC retention time compared with the standard (±2) (Muñoz-Olaya, Matabosch et al., 2008).

### SMase treatment

All experiments (sphingolipid quantification, apoptosis, and TCR nanoclustering) were performed by incubating OT-II WT and CCR5^−/−^ cells (0.2 × 10^6^) with recombinant sphingomyelinase from *Bacillus cereus* (0.5 U/ml; 1 h, 37°C) in serum-free medium. Cells were washed and processed immediately for EM analysis or for sphingolipid quantification as above.

### Quantitative RT-PCR analyses

Total RNA was extracted from human or murine cells using the RNeasy Mini Kit (Qiagen), and cDNA synthesized from 1 µg total RNA (High Capacity cDNA Reverse Transcription Kit, Promega). Quantitative RT-PCR was performed using FluoCycle II SYBR Master Mix (EuroClone) with specific primers (Supplementary Table S1) in an ABI 7300 Real Time PCR System (Applied Biosystems). Results were analyzed using SDS2.4 software.

### CerS2 silencing

Lentiviruses were produced in HEK-293T cells after co-transfection with pGIPZ-shRNA-CerS2 or control plasmids, pSPAX2 and pMD2.G (VSV-G protein) using LipoD293tm (SignaGen). Supernatants were concentrated by ultracentrifugation and supplemented with polybrene (8 µg/ml). Lymphoblasts (3 days post-activation) or 2B4 cells (1.5 × 10^6^ cells/ml) were resuspended in lentiviral supernatant and centrifuged (900 xg, 90 min, 37°C). Transduction efficiency was analyzed after 24 h by FACS. In the case of 2B4 cells, transduced cells were selected with puromycin (2 µg/ml) for 3 days prior to analyses.

### Immunofluorescence analyses

OT-II 10-day WT or CCR5^−/−^ lymphoblasts were plated in poly-L-lysine-coated coverslips (Nunc Lab-Tek Chamber Slide, Thermo Scientific; 50 μg/ml, overnight, 4°C). After adhesion (1 h, 37° C), cells were fixed in 4% PFA (10 min), Triton-X100-permeabilized (0.3% in PBS, 15 min), and blocked with BSA 0.5% in PBS. Samples were incubated (overnight, 4°C) with anti-mouse phospho-GATA1^pSer142^ antibody (1/200), followed by anti-rabbit Ig Alexa-488 secondary antibody (1 h). Coverslips were mounted in Fluoromount-G with DAPI (Southern Biotech); images were acquired with a Zeiss LSM710 and analyzed by a blind observer with NIH Image J software.

### ChIP assay

ChIP assays were performed with the EZ-ChIP Kit (Millipore). In brief, OT-II WT or CCR5^−/−^ lymphoblasts (2 × 10^7^) were fixed (1% PFA, 10 min, RT) and quenched (125 mM glycine, 5 min, RT). Cells were harvested (1 × 10^7^ cells/ml), lysed (15 min, 4°C), and DNA sheared by sonication (45 cycles; 30 s on/off; Bioruptor Pico, Diagenode) in aliquots (0.2 ml). Of each lysate, 1% was stored as input reference, and the remaining material was immunoprecipitated (14 h, 4°C, with rotation) with antibodies to GATA1, histone H3-Lys9, or purified IgG (control). Immune complexes were captured using Protein G magnetic beads (Bio-Rad) and, after washing, eluted with 100 mM NaHCO_3_, 1% SDS; protein/DNA bonds were disrupted with proteinase K (10 µg/µl, 2 h, 62°C). DNA was purified using spin columns, and Cers2 gene promoter sequences analyzed with specific primers (Supplementary Table S1). The relative quantity of amplified product in the input and ChIP samples was calculated (Mira, Carmona-Rodriguez et al., 2018).

### Cell migration assay

OT-II WT or CCR5^−/−^ lymphoblasts (10^6^) were added to the upper chamber of a transwell (3 µm-pore; Corning) and allowed to migrate towards 100 mM CCL4 for 4 h. Migrating cells were quantified by flow cytometry (Cytomics FC500).

### Mathematical and Bayesian analyses

To analyze cluster size distribution, we used a standard χ^2^ test to compare the fraction of clusters of a given size (1, 2, 3, etc.) in each dataset. In all plots, “Random” refers to synthetic distributions of receptors generated randomly.

To quantify the mechanistic relevance of cluster size between random distributions of clusters and clusters in WT and in CCR5^−/−^ CD4^+^ T cells, we used a Bayesian inference model on top of a mechanistic model (Castro et al., 2014). The model assumes that TCR aggregates by incorporating one receptor at a time, with on and off rates that depend on the diffusion properties of the receptor on the membrane, but not existing cluster size. That is,

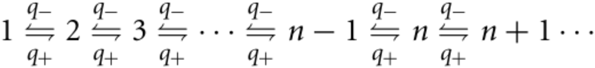

The “affinity” of the process is given by *b* = *q+/q-*, which we also refer to as the clustering or affinity parameter. In the steady state, we can calculate analytically the fraction of clusters of a given size n:

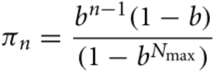

with

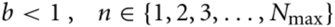

The model was fitted using the Bayesian JAGS code (Kruschke, 2014) (see Supplementary material). The histograms for the number of clusters of a given size *n* (*N_n_*) were modeled as a multinomial distribution with the number of observations, *N*, given by the total count per experiment, and probabilities *π_n_* given by the formulas above. The priors for the clustering parameter *b* are beta distributions with shape parameters *A* and *B* with non-informative uniform priors. Specifically,

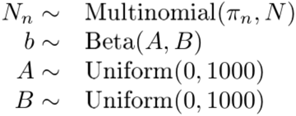

Posterior distribution of the estimated clustering parameter, *b*, is given with the so-called Region of Practical Equivalent (ROPE), defined as the probability of a parameter from a dataset to be explained by another dataset. ROPE quantifies the probability that the observed clustering parameter (and distribution of clusters) in the experiment can be obtained by pure random proximity.

At the molecular level, the kinetic rates q^+^ and q^−^ can be expressed in terms of the diffusion rates of the receptors, kd^+/−^, the receptor size (a), the mean distance between receptors (s), and the correct receptor-receptor binding rates, k^+/−^, through the equations (Lauffenburger & DeLisi, 1983):

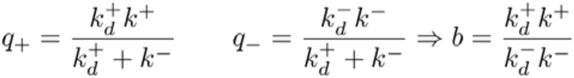

with

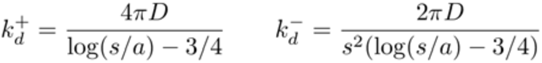

The clustering parameter *b* is independent of the TCR diffusivity (as D is canceled), so the observed TCR nanoclustering differences for WT and CCR5^−/−^ cells would be due to TCR-TCR interactions, as previously reported (Beck-Garcia et al., 2015).

### Identification of transcription factors in CerS promoters

Cers2, Cers3, Cers4, and Cers6 gene coordinates were obtained from the UCSC Genome browser (https://genome.ucsc.edu/), mouse genome version GRCm38/mm10). Known transcription factors for these genes were identified at GTRD v17.04 (http://gtrd17-04.biouml.org/). Venn diagrams were constructed to identify common and specific transcription factors for ceramide synthase genes.

### Statistical analyses

For comparison between two conditions, data were analyzed using parametric Student’s *t*-tests, unpaired, or paired when different treatments were applied to the same sample. Multiple parametric comparisons were analyzed with one-way ANOVA with Bonferroni’s post-hoc test. The χ^2^ test was used to analyze overall distribution of gold particles. All analyses were performed using Prism 6.0 or 7.0 software (GraphPad). Differences were considered significant when p <0.05.

### Data availability

The authors confirm that all relevant data and materials supporting the findings of this study are available on reasonable request. This excludes materials obtained from other researchers, who must provide their consent for transfer.

### Code availability

The Bayesian JAGS code generated in the study is provided as supplementary information.

### Online supplemental material

Figure S1 shows the expression of CCR5 ligands and CCR5 in lymphoblasts, the functionality of CCR5 in these cells, as well as the analysis of TCR nanoclustering in OT-II WT lymphoblasts expanded in IL-2 or IL-15. Fig. S2 shows the comparison of experimental and synthetic TCR nanocluster distribution. Fig. S3 shows a representative FACS analysis of CD4^+^ memory T cells purified by negative selection from OVA/OVA-immunized WT and CCR5^−/−^ mice. Fig. S4 shows the lack of effect of a CCR5 antagonist (TAK-779) on TCR nanoclustering in CCR5-deficient lymphoblasts. Fig. S5 shows that CCR5 deficiency does not affect TCR expression in in vitro-generated lymphoblasts or in vivo isolated memory CD4^+^ T cells. Fig. S6 shows sphingolipid levels in naïve WT and CCR5-deficient OT-II cells, and representative dot plots of WT and CCR5-deficient OT-II cells stained with apoptotic markers. Fig. S7 shows relative mRNA expression levels of enzymes involved in ceramide metabolism. Fig. S8 shows that SMase treatment does not induce apoptosis. Fig. S9 shows the effect of CerS2 silencing in primary CD4^+^ lymphoblasts. Fig. S10 shows that CCR5 blockade with TAK-779 impairs TCR nanoclustering in the CD4^+^ T cell hybridoma 2B4. Fig. S11. shows relative mRNA expression levels of enzymes involved in ceramide metabolism in primary human CD4^+^ lymphoblasts. Table S1 shows the list of primers used in the study. Table S2 shows the Resource Identification Portal accession numbers or catalog numbers of the reagents used in the study. It is also provided as supplemental material the Bayesian code for the R-language used in the study.

## ACKNOWLEDGMENTS

We thank D Sancho and JW Yewdell for rVACV-OVA virus, RM Peregil for technical assistance, MC Moreno and S Escudero for flow cytometry services (CNB), MT Rejas and M Guerra for EM service (CBM-SO), and C Mark for excellent editorial assistance. Fundació ACE would like to thank patients, controls, and the staff who participated in this project. This work was funded by grants from the Spanish Ministerio de Ciencia, Innovación y Universidades (SAF2017–83732-R to SM; FIS2016-78883-C2-2-P to MC; CTQ2017-85378-R; AEI/FEDER, EU), the Instituto de Salud Carlos III (ISCIII) (PI13/02434, PI16/01861 to AR), the Comunidad de Madrid (B2017/BMD-3733; IMMUNOTHERCAN-CM to SM), and the Merck-Salud Foundation (to SM). WWS and CD were supported by the Deutsche Forschungsgemeinschaft (DFG) through BIOSS-EXC294 and CIBSS-EXC 2189 (Project 390939984) and SCHA976/7-1. The Genome Research @ Fundació ACE project (GR@ACE) is supported by the Fundación Bancaria La Caixa, Grifols SA, Fundació ACE, and ISCIII (Ministry of Health, Spain). Fundació ACE is a participating center in the Dementia Genetics Spanish Consortium (DEGESCO).

## AUTHORS’ CONTRIBUTIONS

SM conceived the study. AM-L and RB designed, performed most experiments, and interpreted data. JC and GF performed lipid analyses and CD, liposome experiments. ER-B performed analysis of TCR chains. MC carried out mathematical simulations and Bayesian analyses. MES carried out bioinformatic analyses. IR, LMR, and AR performed *ccr5*Δ32 genotyping and provided samples. WWAS, HMS, and BA contributed ideas and technical support. SM and RB wrote the manuscript. All authors read, discussed and edited the manuscript.

