## Supplemental Figures, Tables and code for "CCR5 deficiency impairs CD4^+^ T cell memory responses and antigenic sensitivity through increased ceramide synthesis"

Supplemental information

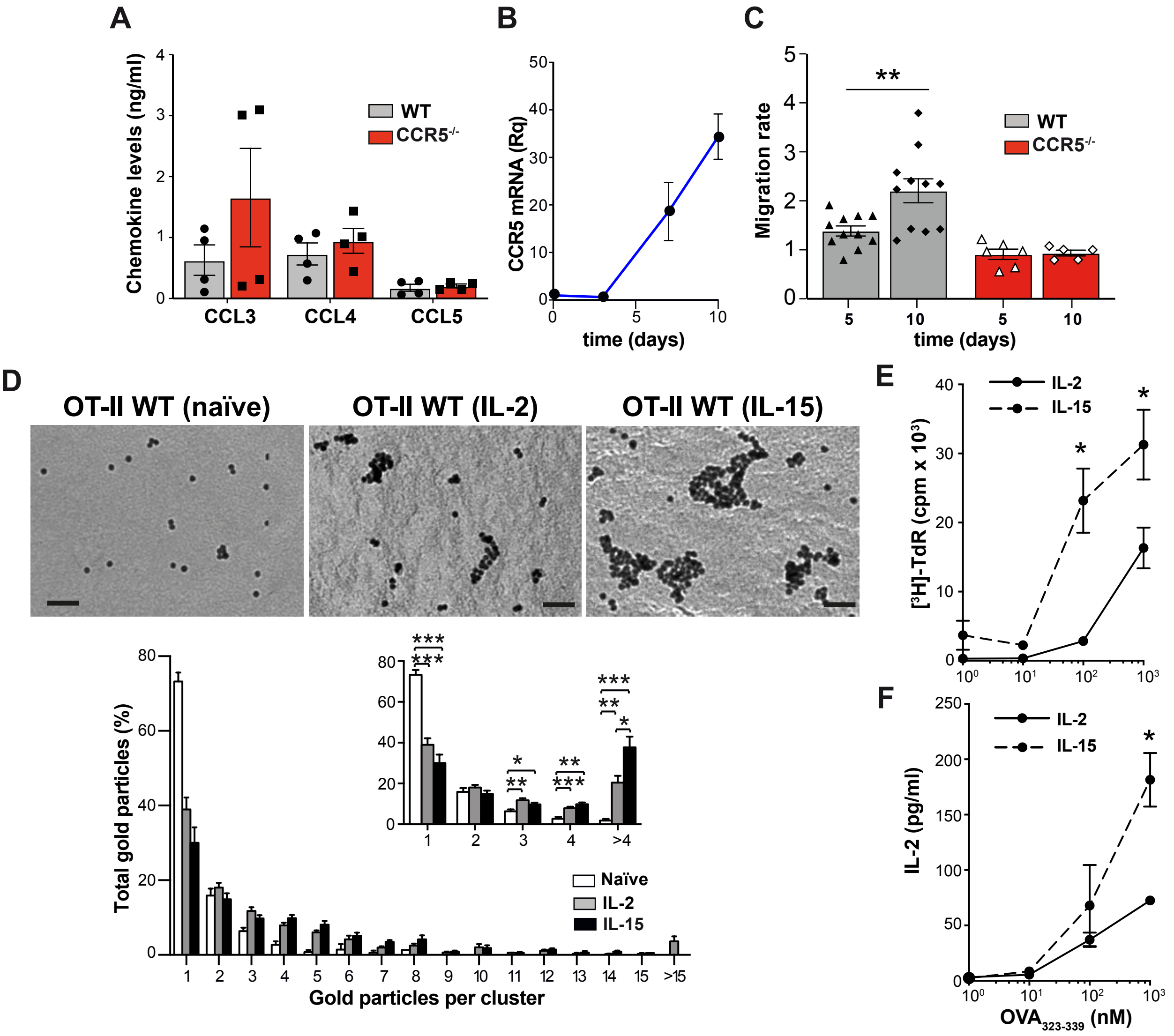


**Figure S1. Characterization of CD4⁺ T lymphoblasts. A.** CCR5 ligand levels in supernatants of day 10 IL-2-expanded OT-II WT and CCR5$^{-/-}$ lymphoblasts. Data shown as mean ± SEM (*n* = 4). **B.** Relative CCR5 mRNA levels in OT-II WT lymphoblasts as in *a*. mRNA was not detected in OT-II CCR5$^{-/-}$ lymphoblasts. **C.** CCL4-induced transmigration of OT-II WT and CCR5$^{-/-}$ lymphoblasts in transwell chambers. Values were normalized to medium without chemoattractant. **D.** Analysis of TCR nanoclustering in OT-II WT naïve cells and lymphoblasts expanded in IL-2 or IL-15. Top, representative small field images showing gold particle distribution in the cell surface replicas of anti-CD3ε-labeled cells. Bottom, quantification (mean ± SEM) of gold particles in clusters of the indicated size in the IL-2- (gray bars; *n* = 6 cells, 27518 particles) and IL-15-expanded lymphoblasts (black; *n* = 8 cells, 27518 particles). Insets show the distribution between clusters of one, two, three, four or more than four particles, and statistical analysis. **E, F.** IL-2- and IL-15-expanded OT-II WT lymphoblasts were restimulated with the indicated concentrations of $OVA_{323-339}$; cell proliferation measured by thymidine incorporation into DNA (*E*) and IL-2 production measured by ELISA (*F*) were determined after 72 h of stimulation. Data shown as mean ± SEM (*n* = 5). * p <0.05, ** p <001, *** p <0.001, two-tailed unpaired Student's *t*-test. Bar, 50 nm.

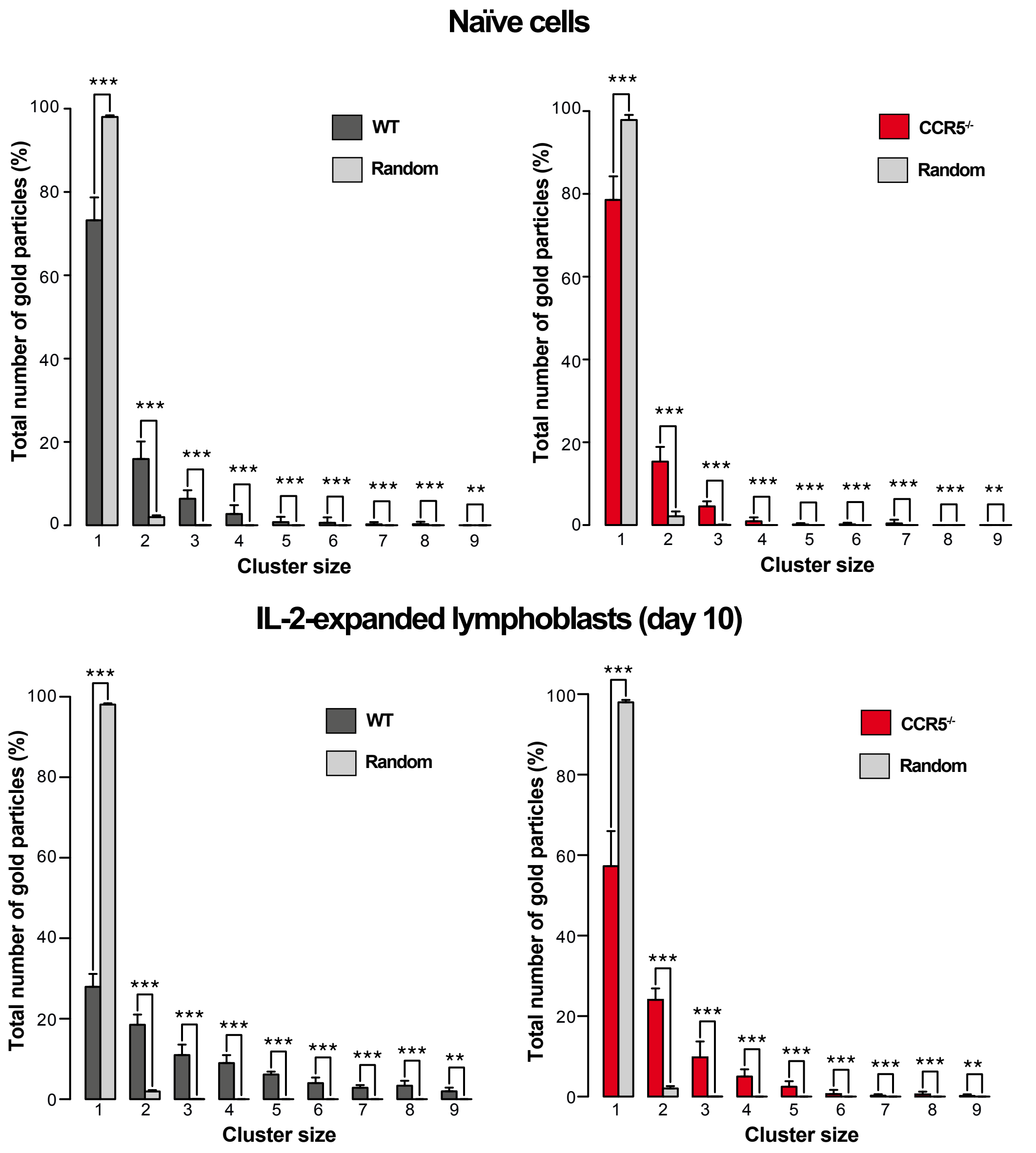


**Figure S2. Comparison of experimental and synthetic TCR multimer distributions.** Percentage of clusters of size *n* (1 to 9) in WT and $CCR5^{-/-}$ OT-II naïve cells and lymphoblasts (day 10) determined experimentally (dark bars), and synthetically random generated receptors (light bars). Student's *t*-test significance for each cluster size is shown above bars (p >0.05 (not significant), * p <0.05, **p <0.01, *** p <0.001). In all cases, the experimental distributions of clusters differ significantly from random proximity between clusters.

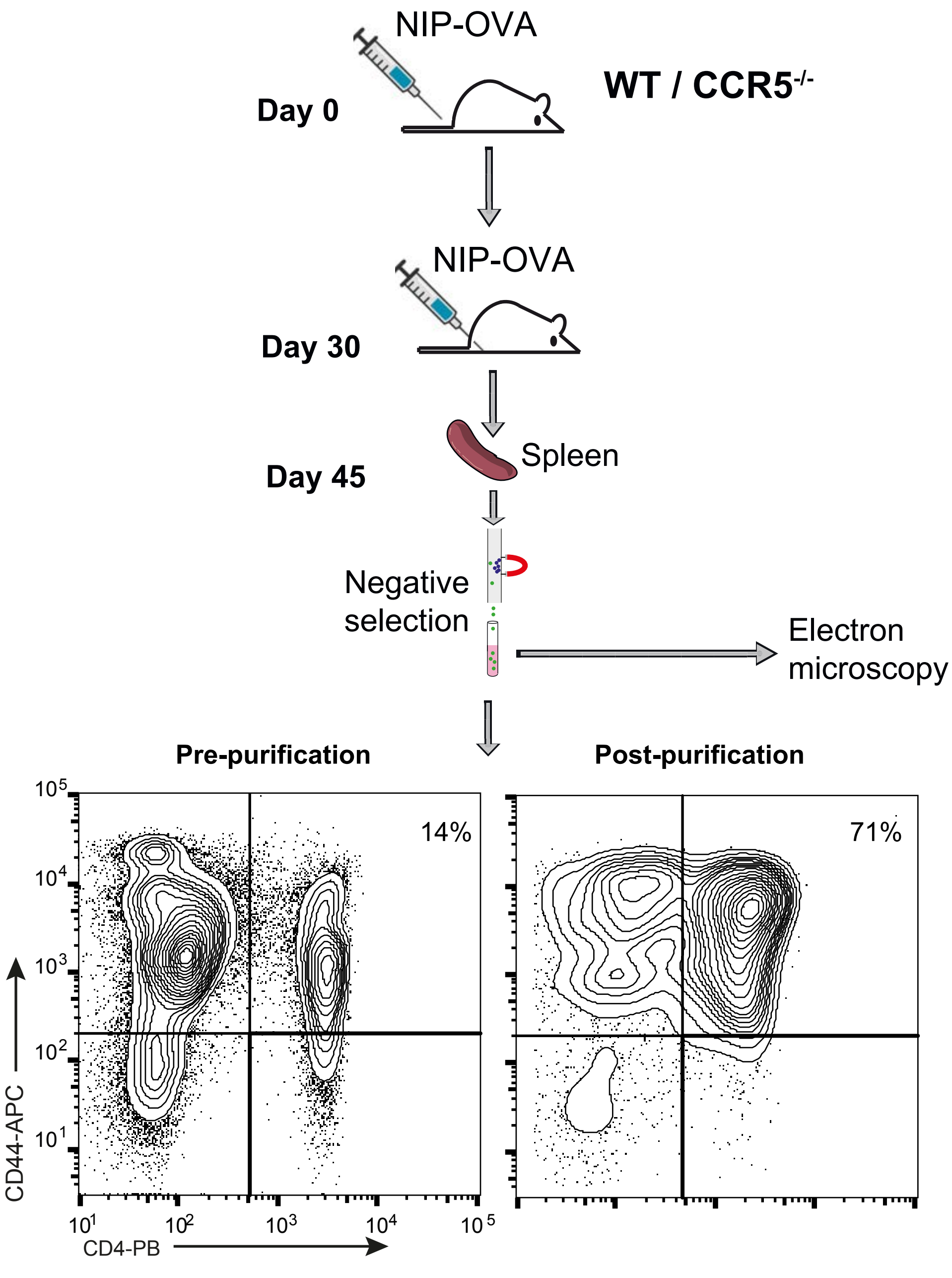


**Figure S3. Characterization of endogenous memory CD4+ T cells uses for electron microscopy studies.** Scheme of the purification of memory cells from OVA/OVA-immunized mice as well as a representative plot showing the characterization of the purified cells by flow cytometry.

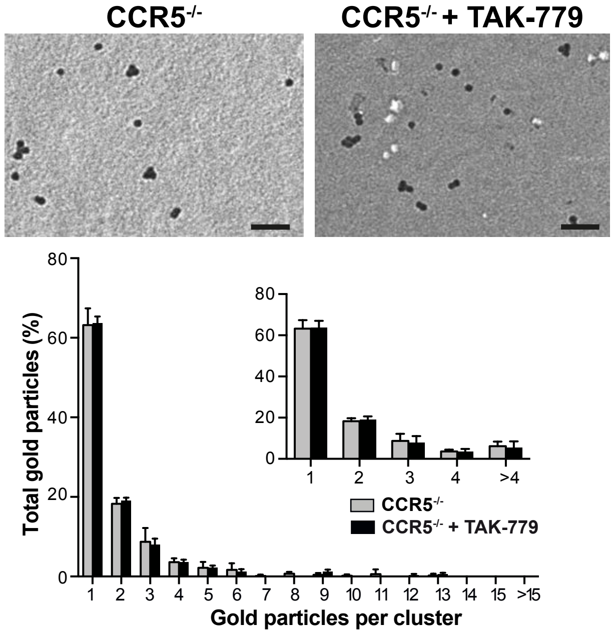


**Figure S4. TAK-779 does not affect TCR nanoclustering in CCR5$^{-/-}$ lymphoblasts.** OT-II CCR5$^{-/-}$ cells were activated with $OVA_{323\text{-}339}$ for 3 days in the presence of TAK-779, and lymphoblasts were generated by expansion with IL-2. Top, representative small field images showing gold particle distribution in cell surface replicas of anti-CD3ε-labeled cells. Bottom, quantification (mean ± SEM) of gold particles in clusters of the indicated size in vehicle-treated (gray bars; $n$ = 6 cells, 5138 particles) and TAK-779-treated lymphoblasts (black; $n$ = 5 cells, 4215 particles). Insets, distribution between clusters of one, two, three, four or more than four particles, and statistical analysis. * $p < 0.05$, one-tailed unpaired Student's $t$-test. Bar, 50 nm.

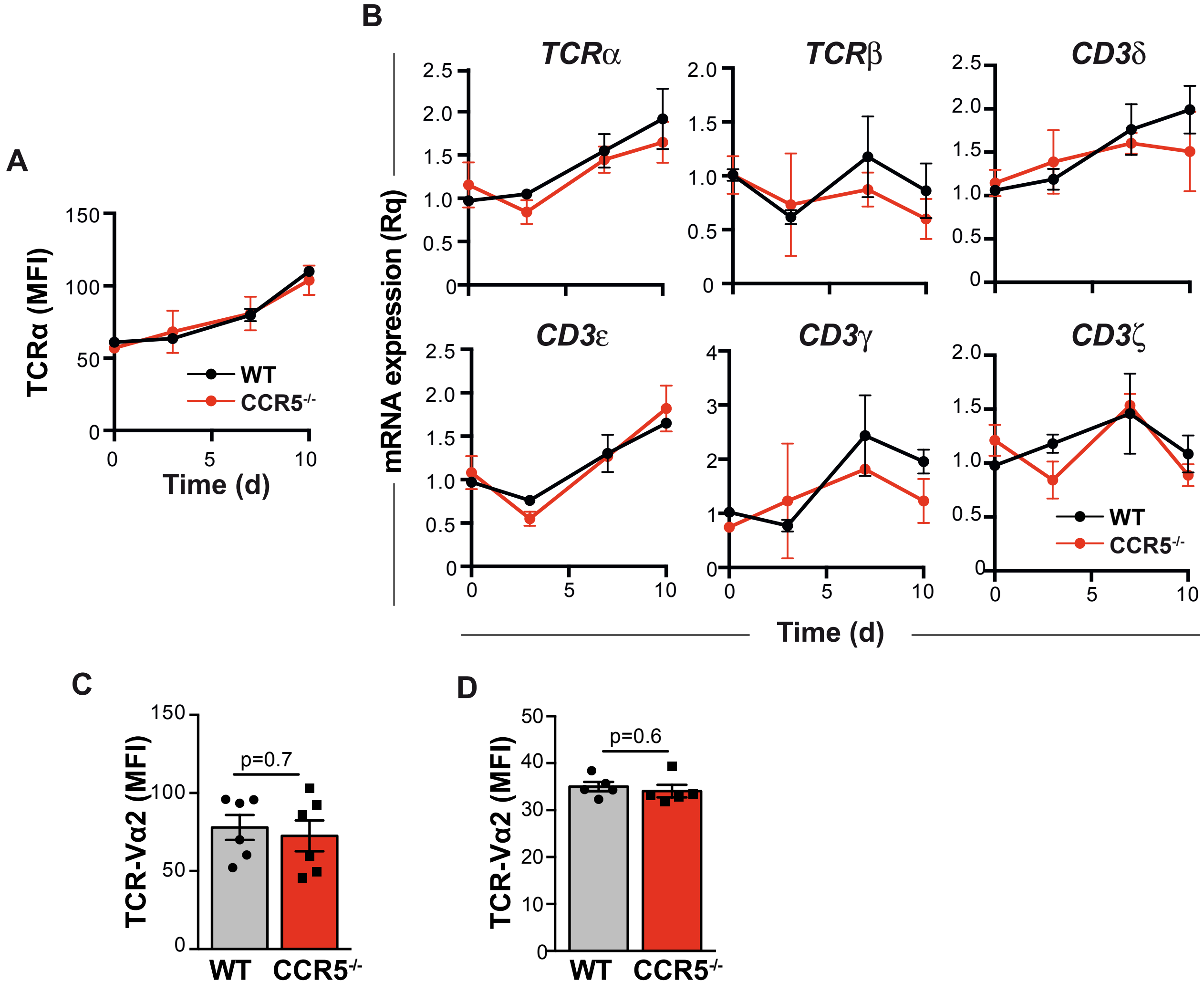


**Figure S5. CCR5 does not affect TCR expression. A.** Quantification of mean fluorescence intensity of TCRα (Vα2) surface staining in IL-2-expanded, $OVA_{323\text{-}339}$-activated WT and $CCR5^{-/-}$ OT-II cells on the days indicated. **B.** Relative mRNA levels for the indicated TCR chains in cells as above. **C.** Mean fluorescence intensity of TCRα (Vα2) surface staining in IL-15-expanded, $OVA_{323\text{-}339}$-activated WT and $CCR5^{-/-}$ OT-II cells. **D.** Mean fluorescence intensity of TCRα (Vα2) surface staining in $CD45.2^{+}/CD4^{+}$ memory cells isolated from NIP-OVA-immunized WT and $CCR5^{-/-}$ mice. In all cases, data shown as mean ± SEM ($n \geq 3$). Differences were not significant using a two-way ANOVA with Bonferroni post-hoc test (*A*, *B*) or two-tailed Student's *t*-test (*C*, *D*).

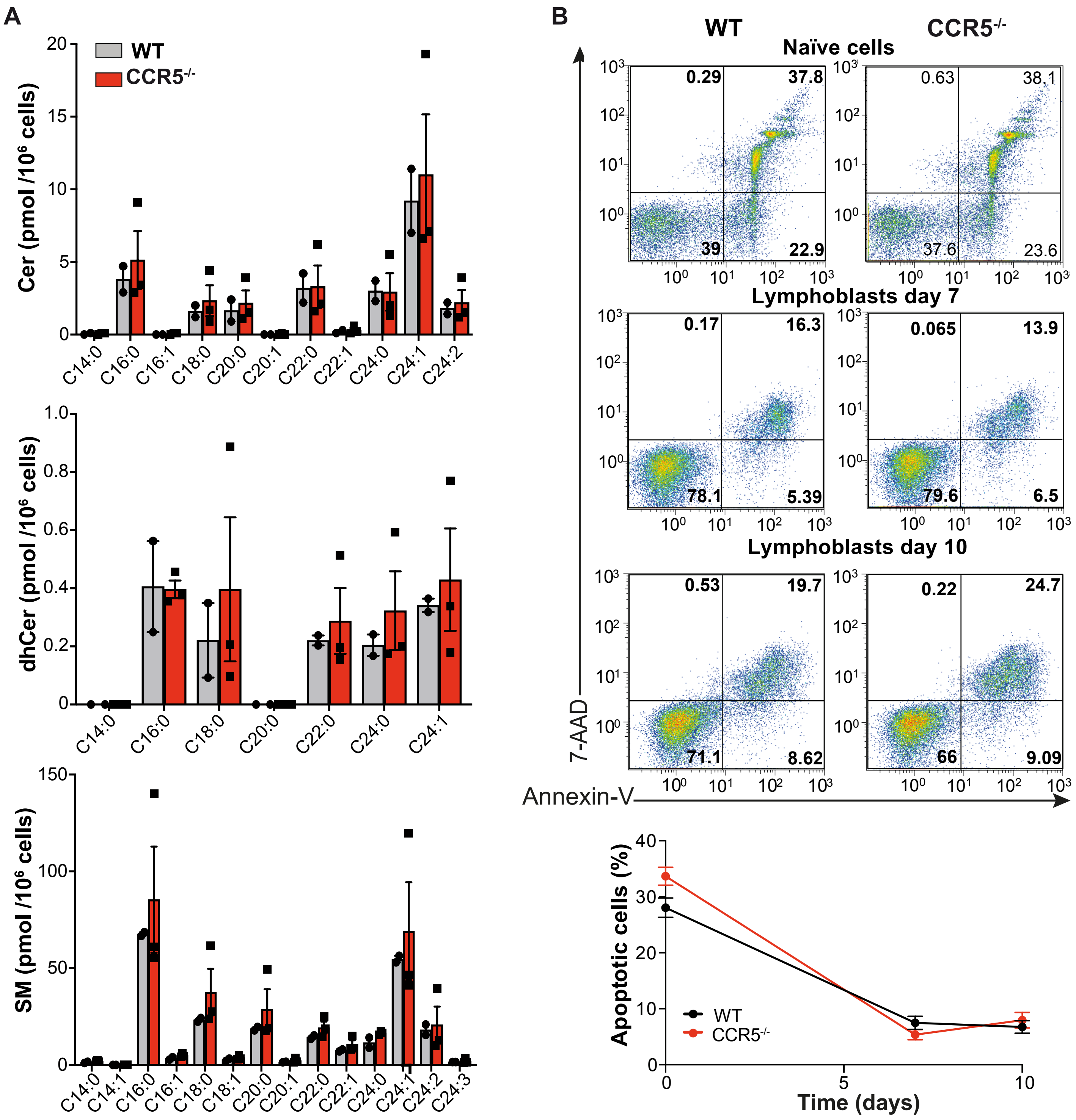


**Figure S6. Analysis of sphingolipids and apoptosis in WT and CCR5$^{-/-}$ naïve OT-II cells. A.** Cer (top), dhCer (center) and SM (bottom) levels in WT and CCR5$^{-/-}$ OT-II naïve cells. Values were normalized to the C17 standards and to cell number in each sample ($n$ = 4). No significant differences were found between genotypes in any lipid species (two-tailed unpaired Student's $t$-test). **B.** Representative dot plots of WT and CCR5$^{-/-}$ OT-II naïve cells and IL-2 lymphoblasts at days 7 and 10, stained for the apoptosis markers 7-aminoactinomycin D (7-AAD) and annexin-V. Numbers represent the percentage of cells. The graph shows the quantification of doubled-stained cells in different experiments (bottom). Data are mean ± SEM ($n$ = 4). No differences were found between genotypes (two-tailed unpaired Student's $t$-test).

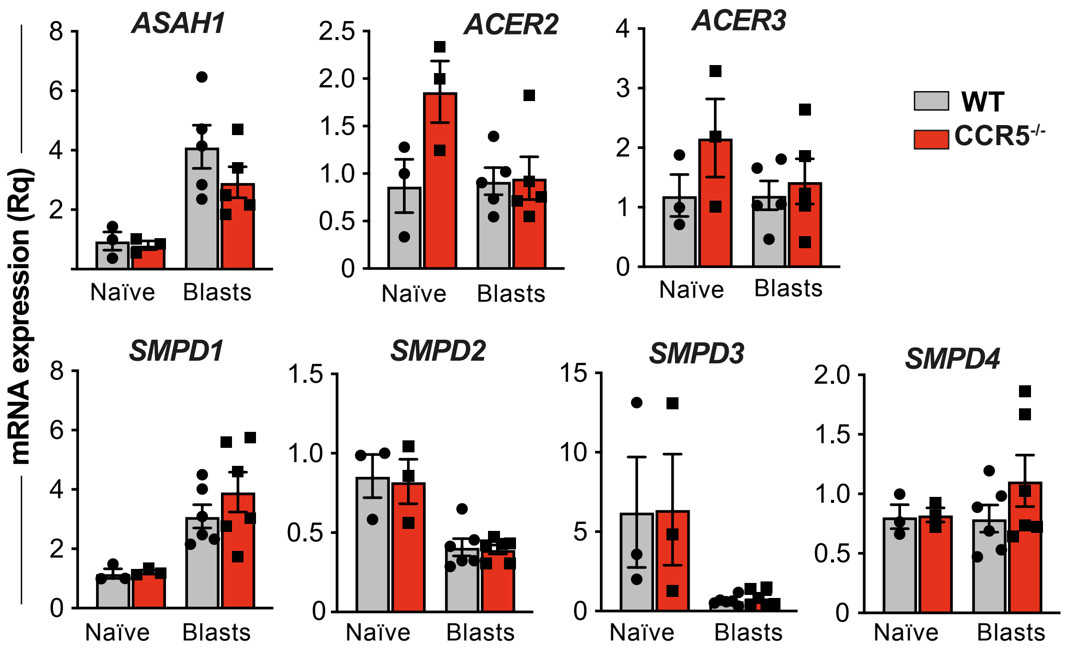


**Figure S7. Characterization of enzymes involved in ceramide metabolism.** Relative mRNA levels for acid ceramidase (ASAH1), alkaline ceramidases 2 and 3 (ACER2, ACER3), acid sphingomyelinase (SMPD1), and neutral sphingomyelinases (SMPD)-2, -3 and -4 in WT and CCR5$^{-/-}$ OT-II naïve cells and lymphoblasts (day 10). Data shown as mean ± SEM ($n$ = 3 or 5). There were no significant differences between genotypes at any given time point (two-tailed unpaired Student's $t$-test).

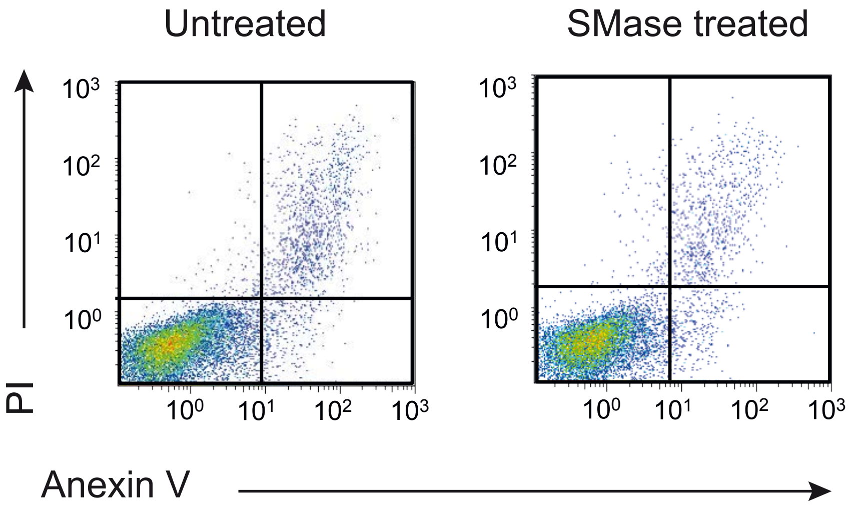


**Figure S8. SMase treatment does not trigger apoptosis in lympholasts.** Day 10 OT-II lymphoblasts were treated with SMase (0.5 U/ml, 1 h, 37ºC) and apoptosis determined by FACS analysis after staining with annexin-V and propidium iodide (PI). Plots shown for untreated (left) and SMase-treated cells (right) in a representative experiment (*n* = 4).

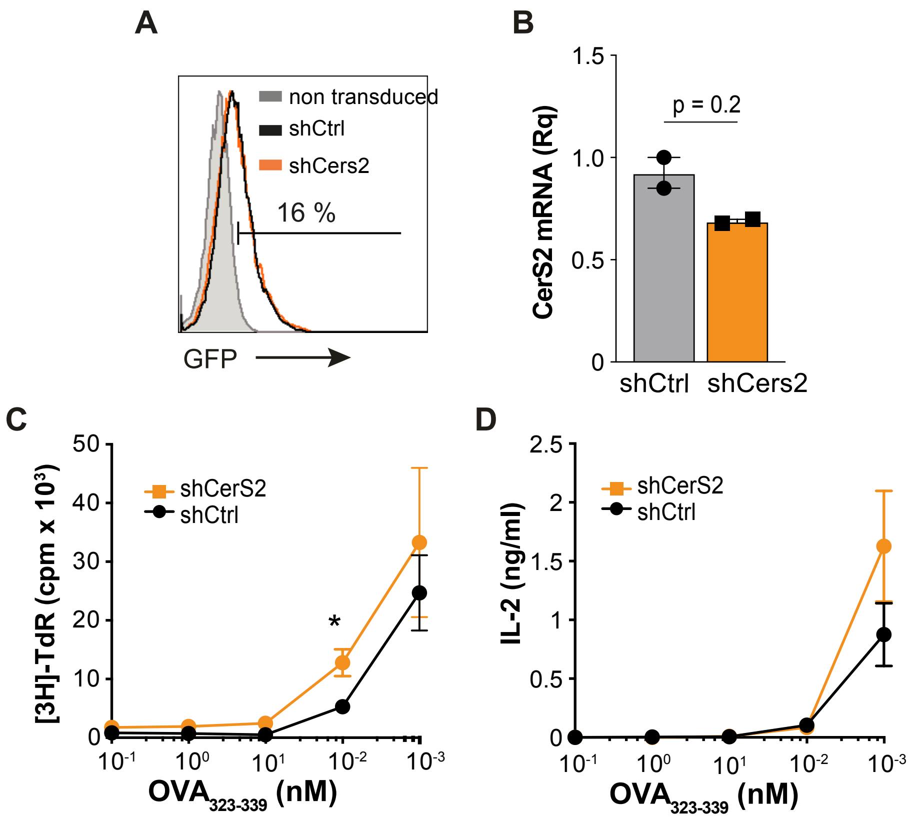


**Figure S9. CerS2 silencing in primary CD4$^+$ lymphoblasts. A.** Histograms showing GFP staining in shCtrl- (black line) and shCerS2-transduced OT-II CCR5$^{-/-}$ lymphoblasts (orange). Data for cells stained at 72 h post-transduction in a representative experiment. Gray area represents staining of non-transduced cells (negative control). The percentage of GFP$^+$ cells is indicated. **B.** Relative CerS2 mRNA levels in cells as in *A*. **C, D.** Determination of thymidine incorporation into DNA (*C*) and IL-2 levels (*D*) in the supernatant of shCtrl- (black line) and shCerS2-transduced (orange line) OT-II CCR5$^{-/-}$ lymphoblasts restimulated for 48 h with $OVA_{323\text{-}339}$ at the indicated concentrations. For *B-D*, data shown as mean ± SEM ($n = 3$). * $p < 0.05$, two-tailed unpaired Student's *t*-test.

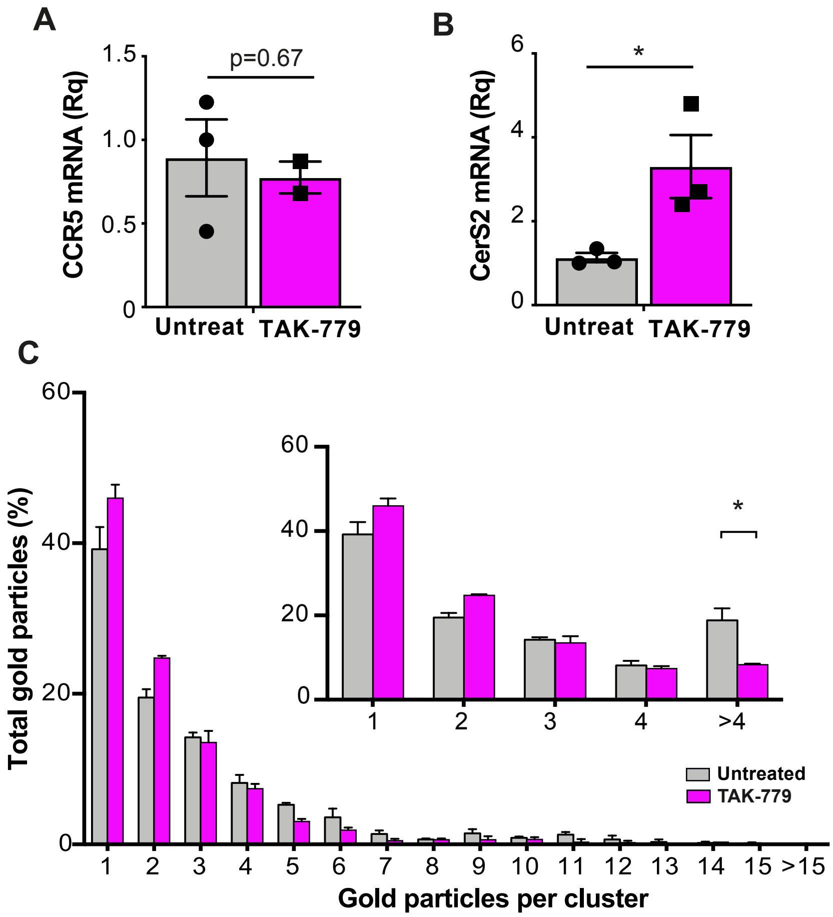


**Figure S10. CCR5 blockade also impairs TCR nanoclustering in the CD4+ T cell hybridoma 2B4. A, B.** Relative mRNA levels of CCR5 (*A*) and CerS2 (*B*) in CD3ε-activated 2B4 cells untreated (gray bars) or TAK-779-treated (10 $\mu$M; pink). **C.** Quantification (mean ± SEM) of gold particles in clusters of the indicated size in cell surface replicas of anti-CD3ε-labeled untreated (gray bars; *n* = 5 cells, 13266 particles) and TAK-779-treated 2B4 cells (black; *n* = 6 cells, 17654 particles). Insets show the distribution between clusters of one, two, three, four or more than four particles, and statistical analysis. * p <0.05, one-tailed unpaired Student's *t*-test.

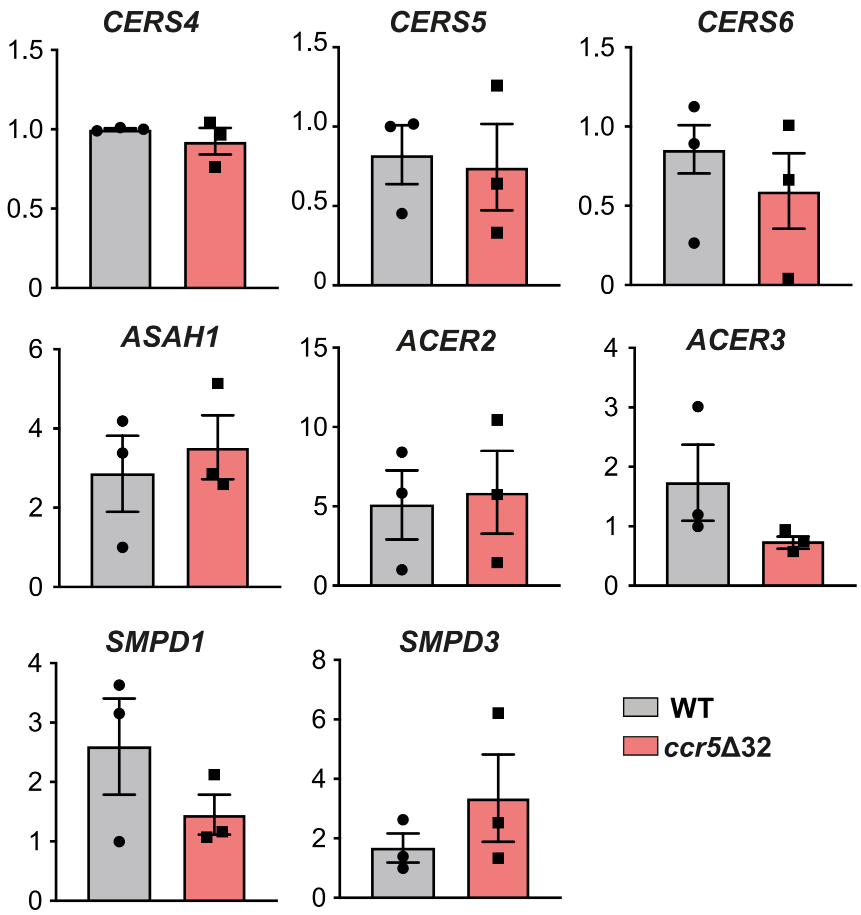


**Figure S11. Characterization of enzymes involved in ceramide metabolism in human CD4⁺ T cells.** Relative mRNA levels for CERS4, CERS5, CERS6, ASAH1, ACER2, ACER3, acid SMase (SMPD1), and neutral SMase (SMPD3) in primary lymphoblasts from healthy donors homozygous for *ccr5*Δ32 (red bars) or who do not bear this polymorphism (WT; gray). Data shown as mean ± SEM ($n$ = 3). There were no significant differences between genotypes for these enzymes (two-tailed unpaired Student's $t$-test).

**Table S1. List of primers used for RT-qPCR analyses**

| Gene Symbol | Primer (5' – 3') Forward | Reverse |
|---|---|---|
| mCerS2 | GGCGCTAGAAGTGGGAAAC | TCGAATGACGAGAAAGAGCA |
| mCerS3 | GCTACACCTCTAGCAAATGCAC | ATCTTTCAACCTGGCGCTCT |
| mCerS4 | AGATAAAGCCCAACCCGGTG | GTCTCCTGAACCAGCGTTGA |
| mCerS5 | CCAATGCTGGTTTCGCCATC | AGAACCAAGGCATCGACCAG |
| mCerS6 | GGAGCTGTCATTTTATTGGTCTTT | GGAACATAATGCCGAAGTCC |
| mASAH1 | TGAAGATGGTGGATCAAAAGC | ACATCTGCAATTCCCCTCA |
| mACER2 | GTGTGGCATATTCTCATCTG | TAAGGGACACCAATAAAAGC |
| mACER3 | TGACCTTGTTCGTCGCTGAG | AGCAATGTACCGCTTCTCCA |
| mSMPD1 | TGGTTCTGGCTCTGTTTGACTCCA | TCAGCTGATCTTGGCGAGACTGTT |
| mSMPD2 | GGTGCTCAACGCCTATGTG | CGTCTGCCTTCTTGGATGTG |
| mSMPD3 | AGAAACCCGGTCCTCGTACT | CCTGACCAGTGCCATTCTTT |
| mSMPD4 | GCCAACGACCTGGACGAGATC | GCGAGTGTGAACTGCCTGAG |
| CCR5 | TCCGTTCCCCCTACAAGAGA | TTGGCAGGGTGCTGACATAC |
| mTcra | CCAGACTGGCAGCAAGAAGAAAAT | TCACAGCTCCCCACCATATTC |
| mTcrb | GTGCTGTGAAGGATGGCAACT | CGGCAGGGTCAGGGTTCT |
| mCD3d | TGTGCAAGTCCATTACCGAAT | AAAGCAGTAGACGCCCAAAG |
| mCD3g | GAGAAGCAAAGAGACTGACATGG | TTATTTGTCTGGGCTACAGTGC |
| mCD3e | AACACTTTCTGGGGCATCCT | ATGTTCTCGGCATCGTCCT |
| mCD3z | GCACGATGGCCTTTACCA | CAAGTGACATCAGCAGGTGAA |
| mGATA-1 | TGCTCTCTTCTTGAGGCATAGATT | CCAGCCCTGCTGTTTAGAGTC |
| mHISTH3 | GCTAAGCTTAACTCTCCCGGT | AAGCGCCCAGCAGCC |
| hCerS2 | GACGGAGTACACGGAGCAG | CGTTCCCACCAGAAGTAATCA |
| hCerS4 | TGGTGCTGCTGTTACACGAT | TGATACTGCATGTAGTTGACCATC |
| hCerS5 | CACATCCTCTCGGTGTTCC | CAGGGTTTGGCAATAAATCG |
| hCerS6 | CGACTGGGTATATTTCCTCTCTG | GGAAGGGTAAGGTCCAACG |
| hASAH1 | CACGCTGATTGGGTGTGTA | CGATGTTCAGCTTGTATTTCTTGA |
| hACER2 | TGTGGTTCCCCAGAAGGTAT | ACGTCGTAACCGCAGACAG |
| hACER3 | CCTGAGATATAGGCCAAAAGTGA | GCCAGGTCTATGGTAGGTGCT |
| hSMPD1 | TGGCTCTATGAAGCGATGG | TGGGGAAAGAGCATAGAACC |
| hSMPD3 | TGGTACCCAAGAACTGCTACG | AAAACCCAGAAACTGCCTTG |
| 18S rRNA | GAGAAACGGCTACCACATCC | GGGTCGGGAGTGGGTAAT |

### Table S2. RRID accession number and catalog reference for the reagents used

| REAGENT | SOURCE | IDENTIFIER |
|---|---|---|
| **Antibodies** | | |
| Anti-mouse $V\alpha_2$TCR-PE (B20.1) | BD-Biosciences | Cat# 553289, RRID:AB 394760 |
| Anti-mouse CD25-PE (PC61) | BD-Biosciences | Cat# 553866, RRID:AB 395101 |
| Anti-mouse CD45.2-FITC (104) | BD-Biosciences | Cat# 561874,RRID:AB 10894189 |
| Anti-mouse CD62L-FITC (MEL-14) | BD-Biosciences | Cat# 553150,RRID:AB 394665 |
| Anti-mouse CD62L-APC (MEL-14) | BD-Biosciences | Cat# 553152,RRID:AB 398533 |
| Anti-mouse CD69-PeCy7 (H1.2F3) | BD-Biosciences | Cat# 552879,RRID:AB 394508 |
| Anti-mouse CXCR5 biotinylated (2G8) | BD-Biosciences | Cat# 551960,RRID:AB 394301 |
| Anti-mouse biotin CD3ε (145-2C11) | BD-Biosciences | Cat# 553239,RRID:AB 394728 |
| Anti-mouse unlabeled CD3ε (145-2C11) | BD-Biosciences | Cat# 553057,RRID:AB_394590 |
| Anti-human biotin CD3 (OKT3) | Thermo Fisher Scientific | Cat# 13-0037-80, RRID:AB_1234956 |
| Anti-mouse CD4-PeCy7 (RM4.5) | Thermo Fisher Scientific | Cat# 25-0042-81, RRID:AB_469577 |
| Anti-mouse CD4-efluor450 (RM4.5) | Thermo Fisher Scientific | Cat# 48-0042-80, RRID:AB_1272231 |
| Anti-mouse CD4-PacificBlue (RM4.5) | Thermo Fisher Scientific | Cat# MCD0428, RRID:AB_10372505 |
| Anti-mouse IFNγ-APC (XMG1.2) | Thermo Fisher Scientific | Cat# 17-7311-82, RRID:AB_469504 |
| Anti-mouse PD1-efluor780 (J43) | Thermo Fisher Scientific | Cat# 47-9985-82, RRID:AB_2574002 |
| Anti-mouse phospho-GATA1-pSer142 | Thermo Fisher Scientific | Cat# PA5-37581, RRID:AB_2554189 |
| Anti-mouse CD44-Pe/Cy5 (IM7) | BioLegend | Cat# 103009, RRID:AB_312960 |
| Anti-mouse CD44-APC (IM7) | BioLegend | Cat# 103011, RRID:AB_312962 |
| Anti-mouse CerS2 (1A6) | Novus Biologicals | Cat# H00029956-M01A, RRID:AB_2132954 |
| Anti-mouse CerS3 (6C12) | Novus Biologicals | Cat# H00204219-M02 |
| Anti-mouse CerS6 | Novus Biologicals | Cat# H00253782-M01, RRID:AB_2133107 |
| Anti-mouse CerS4 (polyclonal) | Sigma-Aldrich | Cat# SAB4503164, RRID:AB_10746317 |
| Anti-mouse CD3ζ (449) | Purified from hybridoma, this study | N/A |
| Anti-mouse β-actin (AC-15) | Sigma-Aldrich | Cat# A1978, RRID:AB_476692 |
| Anti-mouse GATA1 (ab11852) | Abcam | Cat# ab11852, RRID:AB_298635 |
| Anti-acetyl histone H3Lys9 (CS200583) | EMD-Millipore | Cat# 07-352, RRID:AB_310544 |
| purified IgG rabbit (PP64) | EMD-Millipore | Cat# PP64, RRID:AB_97852 |
| **Virus strains** | | |
| rVACV-OVA virus | a gift of J.W.Yewdell; NIH, Bethesda, MD | N/A |
| **Biological Samples** | | |
| Healthy adult blood samples | Fundació ACE (Barcelona, Spain) | NA |
| **Chemicals, peptides, recombinant proteins** | | |
| OVA(323–339) peptide | CNB Peptide facility | N/A |
| Sphingomyelinase (Bacillus cereus) | Sigma-Aldrich | Cat# S7651 |
| TAK-779 | Sigma-Aldrich | Cat# SML0911 |

| | | |
|---|---|---|
| AMD-3100 | Sigma-Aldrich | Cat# 239820 |
| poly-L-lysine | Sigma-Aldrich | Cat# P4707 |
| Ceramide (bovine spinal cord) | Sigma-Aldrich | Cat# 22244 |
| Protein A-Gold | Sigma Aldrich | Cat# P6730 |
| Streptavidin-agarose | Sigma Aldrich | Cat# S1638 |
| Ceramide from bovine spinal cord | Sigma Aldrich | Cat# 22244 |
| NIP-OVA | Biosearch Technologies | Cat# N-5041 |
| NIP-KLH | Biosearch Technologies | Cat# N-5042 |
| Aluminium hydroxide gel | InvivoGen | Cat# vac-alu-250 |
| Streptavidin APC | Thermo Fisher Scientific | Cat# 17-4317-82 |
| Streptavidin PerCP-Cy5 | Thermo Fisher Scientific | Cat# 45-4317-82; RRID: AB_10311495 |
| Dynabeads™ M-450 Tosylactivated | Thermo Fisher Scientific | Cat# 14013 |
| Proteinase K | Thermo Fisher Scientific | Cat# 25530049 |
| Streptavidin BV786 | BD Bioscience | Cat# 563858 |
| Recombinant murine CCL4 | PeproTech | Cat# 250-32 |
| Recombinant murine IL-2 | PeproTech | Cat# 212-12 |
| Recombinant murine IL-15 | PeproTech | Cat# 210-15 |
| Recombinant human IL-2 | PeproTech | Cat# AF-200-02 |
| LIVE/DEAD™ Fixable Near-IR Dead Cell Stain Kit | Molecular Probes | Cat# 10154363 |
| DAPI Fluoromount-G | Southern Biotech | Cat# 0100-20 |
| Thymidine, [Methyl-3H] | Pelkin Elmer | Cat# NET027W001MC |
| SureBeads™ Protein G Magnetic Beads | Bio Rad | Cat# 1614023 |
| LipoD293™ In Vitro DNA Transfection Reagent | SignaGen Laboratories | Cat# SL100668 |
| Cholesterol | Avanti Polar Lipids | Cat#700100 |
| Soybean L-α-phosphatidylcholine | Avanti Polar Lipids | Cat#840054C |
| Egg Sphingomyelin | Avanti Polar Lipids | Cat#860061C |
| C12 Ceramide (d18:1/12:0) | Avanti Polar Lipids | Cat# 860512 |
| C16 Ceramide (d18:1/16:0) | Avanti Polar Lipids | Cat# 860516 |
| C18 Ceramide (d18:1/18:0) | Avanti Polar Lipids | Cat# 860518 |
| C24 Ceramide (d18:1/24:0) | Avanti Polar Lipids | Cat# 860524 |
| C24:1 Ceramide (d18:1/24:1(15Z)) | Avanti Polar Lipids | Cat# 860525 |
| C16 Dihydroceramide (d18:0/16:0) | Avanti Polar Lipids | Cat# 860634 |
| C18 Dihydroceramide (d18:0/18:0) | Avanti Polar Lipids | Cat# 860627 |
| C24 Dihydroceramide (d18:0/24:0) | Avanti Polar Lipids | Cat# 860628 |
| C24:1 Dihydroceramide (d18:0/24:1(15Z)) | Avanti Polar Lipids | Cat# 860629 |
| 12:0 SM (d18:1/12:0) | Avanti Polar Lipids | Cat# 860583 |
| 16:0 SM (d18:1/16:0) | Avanti Polar Lipids | Cat# 860584 |
| 18:0 SM (d18:1/18:0) | Avanti Polar Lipids | Cat# 860586 |
| 24:0 SM | Avanti Polar Lipids | Cat# 860592 |
| 24:1 SM | Avanti Polar Lipids | Cat# 860593 |
| C12 Glucosyl(β) Ceramide (d18:1/12:0) | Avanti Polar Lipids | Cat# 860543 |
| C17 sphinganine (d17:0) | Avanti Polar Lipids | Cat# 860654 |
| C17 sphinganine-1-phosphate (d17:0) | Avanti Polar Lipids | Cat# 860655 |
| Chloroform | JT Baker | Cat# 15588534 |
| Water for LC-MS | JT Baker | Cat# 15568664 |
| Methanol for LC-MS | Fisher Chemical | Cat# 15611630 |
| Ammonium formate for LC-MS | Fisher Chemical | Cat# 11377490 |
| Formic Acid for LC-MS | Fluka | Cat# 15671400 |

| | | |
|---|---|---|
| **Critical commercial assays** | | |
| ELISA MAX™ Deluxe Set Mouse IL-2 | BioLegend | Cat# 431004 |
| Mouse Memory T cell CD4+/CD62L-/CD44hi Column Kit | R&D Systems | Cat# MCD45 |
| Dynabeads™ Untouched™ Mouse CD4 Cells Kit | ThermoFisher | Cat# 11415D |
| Amplex™ Red Cholesterol Assay Kit | ThermoFisher | Cat# A12216 |
| EasySep™ Human CD4+ T Cell Enrichment Kit | StemCell | Cat# 19052 |
| SBA Clonotyping System-HRP | SouthernBiotech | Cat# 5300-05 |
| RNeasy Mini Kit | QIAGEN | Cat# 74104 |
| EZ-ChIP™ | Millipore | Cat# 17-371 |
| **Experimental models: Cell lines** | | |
| HEK-293 T | ATCC | Cat# CRL-3216, RRID:CVCL_0063 |
| 2B4 T cell hybridoma | J. Ashwell, Bethesda | RRID:CVCL_4Z38 |
| M.mζ-SBP | (Swamy and Schamel, 2009) | N/A |
| **Experimental models: Organisms** | | |
| B6.129P2-Ccr5tm1Kuz (CCR5$^{-/-}$) | The Jackson Laboratory | Cat# JAX:005427, RRID:IMSR_JAX:005427 |
| C57BL/6J | The Jackson Laboratory | Cat# JAX:000664, RRID:IMSR_JAX:000664 |
| B6.Cg-Tg(TcraTcrb)425Cbn/J (OT-II) | The Jackson Laboratory | Cat# JAX:004194, RRID:IMSR_JAX:004194) |
| OT-II-CCR5$^{-/-}$ | (González-Martín et al., 2011) | N/A |
| B6.SJL-Ptprca Pepcb/Boy | The Jackson Laboratory | Cat# JAX:002014, RRID:IMSR_JAX:002014 |
| CD3ε$^{-/-}$ | (DeJarnette et al., 1998) | N/A |
| **Oligonucleotides** | | |
| Primers for qRT-PCR, see Table S1 | | N/A |
| **Software and Algorithms** | | |
| Bayesian JAGS code | | Provided as supplementary material |
| FlowJo v10 | Tree Star | RRID:SCR_008520 |
| GraphPad Prism v6 & v7 | GraphPad | RRID:SCR_002798 |
| NIH Image J | NIH Image | RRID:SCR_003073 |
| UCSC Genome browser | University of California Santa Cruz | RRID:SCR_005780 |
| GTRD v17.04 | Gene Transcription Regulation Database | http://gtrd17-04.biouml.org/ |
| Venny 2.1 | BioinfoGP, CNB/CSIC | RRID:SCR_016561 |
| Adobe Photoshop CS5 | Adobe Software | RRID:SCR_014199 |
| Adobe Illustrator CS5 | Adobe Software | RRID:SCR_010279 |
| **Others** | | |
| Brefeldin A | Sigma-Aldrich | Cat# B7651 |
| IntraPep Permeabilization reagent | Beckman Coulter | Cat# A07803 |
| GIPZ lentiviral mouse CerS2 shRNA (clones V3LMM_454307, V3LMM_454309, V3LMM_454311) | Dharmacon | Cat# RMM4532-EG76893 |

| | | |
|---|---|---|
| Bio-Beads™ SM-2 Resin | BioRad | Cat# 1523920 |
| Acquity C8 UPLC column | Waters | Cat# 186002878 |

**Bayesian code for the R-language**

```
# require(rjags)
# The array "p" contains 4 different cluster size counts from different
experiments for the same mouse and time
cluster.jags.multix4 <- function(p) {
  p1 <- p[1,]
  p2 <- p[2,]
  p3 <- p[3,]
  p4 <- p[4,]
  data <-
list(p1=p1,p2=p2,p3=p3,p4=p4,m=length(p1),N1=sum(p1),N2=sum(p2),N3=sum(p3),N4=su
m(p4))

  modelstring="
  model {
  for(n in 1:(m-1)) {
     pi1[n] <- b1^(n-1)*(1-b1) # Analytical distribution described in the main
text
     pi2[n] <- b2^(n-1)*(1-b2)
     pi3[n] <- b3^(n-1)*(1-b3)
     pi4[n] <- b4^(n-1)*(1-b4)
  }
  pi1[m] <- b1^(m-1)  # Analytical distribution described in the main text
  pi2[m] <- b2^(m-1)
  pi3[m] <- b3^(m-1)
  pi4[m] <- b4^(m-1)

  p1 ~ dmulti(pi1,N1) # The counts are given by a multinomial distribution with
probabilities "pi"
  p2 ~ dmulti(pi2,N2)
  p3 ~ dmulti(pi3,N3)
  p4 ~ dmulti(pi4,N4)

  b1 ~ dbeta(A,B) # Priors for the parameters b1.
  b2 ~ dbeta(A,B)
  b3 ~ dbeta(A,B)
  b4 ~ dbeta(A,B)
  A ~ dunif(0,1000) # Hyperpriors for A and B (0 = uniform distribution,
Infinity=peaked distribution)
  B ~ dunif(0,1000)
  }"

  model=jags.model(textConnection(modelstring), data=data,n.chains = 3) # Create
jags model
  update(model,n.iter=10000) # Burning phase of the MCMC model
  output=coda.samples(model=model,variable.names=c("A","B","b1","b2","b3","b4"),
n.iter=15000, thin=1) # Sample data
  print(summary(output)) # Print estimated parameters
  return(output) # return matrix of results for post-processing
}
```
